# LABRAT reveals association of alternative polyadenylation with transcript localization, RNA binding protein expression, transcription speed, and cancer survival

**DOI:** 10.1101/2020.10.05.326702

**Authors:** Raeann Goering, Krysta L. Engel, Austin E. Gillen, Nova Fong, David L. Bentley, J. Matthew Taliaferro

## Abstract

The sequence content of the 3′ UTRs of many mRNA transcripts is regulated through alternative polyadenylation (APA). The study of this process using RNAseq data, though, has been historically challenging. To combat this problem, we developed LABRAT, an APA quantification method. LABRAT takes advantage of newly developed transcriptome quantification techniques to accurately determine relative APA site usage and how it varies across conditions. Using LABRAT, we found consistent relationships between gene-distal APA and subcellular RNA localization in multiple cell types. We also observed connections between transcription speed and APA site choice as well as tumor-specific transcriptome-wide shifts in APA in hundreds of patient-derived tumor samples that were associated with patient prognosis. We investigated the effects of APA on transcript expression and found a weak overall relationship, although many individual genes showed strong correlations between APA and expression. We interrogated the roles of 191 RNA-binding proteins in the regulation of APA, finding that dozens promote broad, directional shifts in relative APA isoform abundance both *in vitro* and in patient-derived samples. Finally, we find that APA site shifts in the two classes of APA, tandem UTRs and alternative last exons, are strongly correlated across many contexts, suggesting that they are coregulated.

## INTRODUCTION

During the co-transcriptional processing of a pre-mRNA, the 3′ end of the transcript is cleaved and a polyadenine tail is added that promotes the stability and translation of the resulting message (Beilharz and Preiss, 2007; Shi et al., 2009). The site where this cleavage occurs determines the sequence content of the 3′ UTR of the transcript. Regulatory cis-element sequences can therefore be either included or excluded from the 3′ UTR of the transcript through modulation of where the cleavage and polyadenylation event happens. This regulation of transcript sequence content through alternative polyadenylation (APA) occurs in the majority of genes in yeast, plant, and mammalian genomes (Derti et al., 2012; Ozsolak et al., 2010; Sherstnev et al., 2012; Wu et al., 2011).

The cleavage and polyadenylation reaction is performed by the core CSTF and CPSF complexes and CFIm which associate with RNA polymerase II (Pol II) transcription complexes (Glover-Cutter et al., 2008; Venkataraman et al., 2005) and together recognize specific sequence elements within 3′ UTRs to determine sites of 3′ end processing (Tian and Manley, 2017). The abundance of these general CPA factors as well as several other RBPs have been found to regulate the relative usage of alternative polyadenylation sites within a transcript (Gruber et al., 2012; Li et al., 2015; Martin et al., 2012; Masamha et al., 2014; Takagaki et al., 1996; Zhu et al., 2018).

Regulation by these factors results in the large variation in 3′ UTR content seen across tissues and developmental stages (Lianoglou et al., 2013). Specific tissues, most notably neuronal tissues, are associated with preferential use of gene-distal or downstream APA sites (Miura et al., 2013). Similarly, the broad use of gene-proximal or distal APA sites can be developmentally regulated. Undifferentiated, proliferating cells generally display enriched usage of proximal APA sites while more differentiated cells show shifts towards increased usage of distal APA sites (Ji et al., 2009; Sandberg et al., 2008). This phenomenon has also been connected to cancer progression where increased usage of proximal APA sites in key oncogenes was associated with elevated cell proliferation and oncogenic transformation (Mayr and Bartel, 2009; Sandberg et al., 2008).

Alternative polyadenylation exists in two structurally distinct forms. The first, which we will refer to as “tandem UTRs” occurs when multiple APA sites are found within the same terminal exon (**Figure 1B, top**). The second, which we will refer to as “alternative last exons” or “ALEs”, occurs when multiple APA sites are found within different terminal exons (**Figure 1B, bottom**). Regulation of the choice between alternative tandem UTRs can be viewed as a competition between a proximal upstream poly(A) site that is transcribed first with a distal downstream site that is transcribed second. Similarly the choice between ALE’s can be viewed as a competition between recognition of a proximal 5’ splice site and a distal poly(A) site (Peterson and Perry, 1989). It is not known whether the two forms of APA are subject to common regulatory mechanisms but in this regard it is interesting to note that transcription speed has been reported to influence the competition between alternative splice sites and tandem poly(A) sites (Liu et al., 2017).

**Figure 1.**
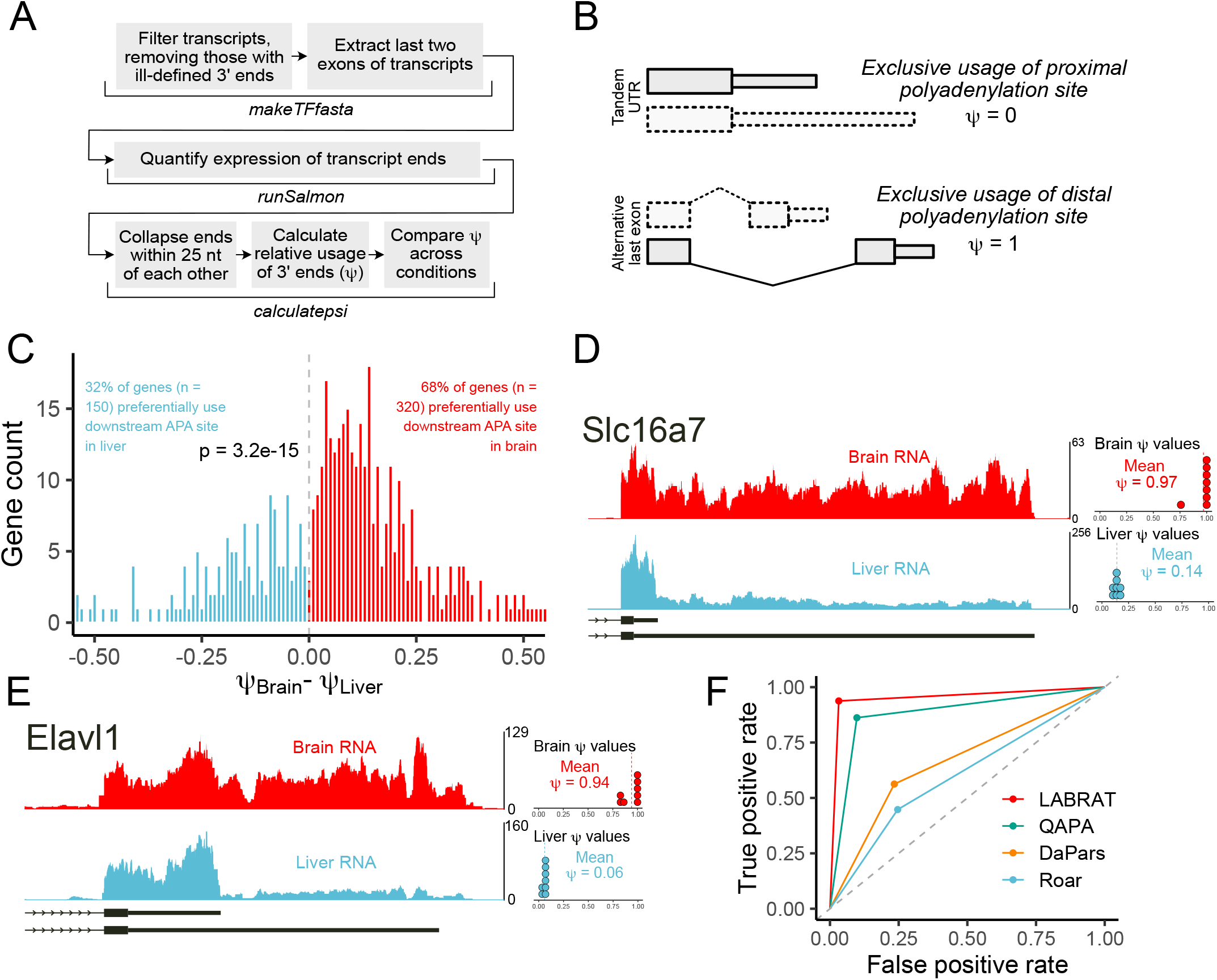
Quantifying changes in alternative polyadenylation with LABRAT. (A) LABRAT computational pipeline. (B) Explanation of *ψ* as a metric of polyadenylation site choice. Genes that exclusively use upstream or gene-proximal sites have *ψ* values of 0 while those that exclusively use downstream or gene-distal sites have *ψ* values of 1. The two transcript structures associated with alternative polyadenylation, tandem UTRs and alternative last exons, are diagrammed. (C) Comparison of *ψ* values in mouse brain and liver RNA for genes whose *ψ* value was significantly different between these tissues. (D-E) RNA coverage profiles of two genes with differential polyadenylation site usage in mouse brain and liver tissues. Dots represent *ψ* values calculated in each of 8 replicates. (F) Benchmarking of LABRAT performance against other widely used software package for quantification of alternative polyadenylation from RNAseq data.

The study of APA using high-throughput RNA sequencing has been facilitated through a handful of software packages aimed at quantifying changes in relative APA site usage across conditions (Grassi et al., 2016; Ha et al., 2018; Xia et al., 2014). However, quantifying APA from transcriptomic alignments can be difficult. Due to their shared isoform structure, different APA isoforms often contain a considerable amount of sequence in common. If the APA quantification software relies on these transcriptomic alignments (Grassi et al., 2016; Xia et al., 2014), this can make assigning reads to a specific isoform challenging. Newer transcriptome quantification techniques that assign reads to transcripts by comparing their sequence contents are better equipped to handle this problem. (Bray et al., 2016; Ha et al., 2018; Patro et al., 2017).

To take advantage of this advance in isoform quantification and apply it to the analysis of APA, we developed LABRAT (Lightweight Alignment-Based Reckoning of Alternative Three-prime ends). A particular advantage of our approach is that it permits rapid analysis of large numbers of publically available RNA-seq data sets including patient samples. Here, we applied this approach to tens of thousands of RNAseq samples to study processes and factors that regulate APA as well as the consequences of APA site choice on transcript fate.

## RESULTS

### Quantification of alternative polyadenylation with LABRAT

To quantify relative alternative polyadenylation site usage from RNAseq data, LABRAT takes a genome annotation file and first searches the annotation for tags that define transcripts with ill-defined 3′ ends in order to filter and remove them from further analysis (**Figure 1A**). Because annotations are isoform-based, they are often rigid in their explicit connection of upstream alternative splicing events to downstream APA sites, even though this connection may not be accurate. Therefore, to exclude spurious contributions of upstream alternative splicing events to APA site quantification, we extracted the final two exons of every transcript and the expression of these transcript “terminal fragments” was quantified using Salmon (Patro et al., 2017).

For each gene, alternative polyadenylation sites are then defined using terminal fragments. Terminal fragments with 3′ ends within 25 nt of other 3′ ends are grouped together to define a single polyadenylation site, and the sites are ordered from most gene-proximal to most gene-distal. Each APA site within a gene is assigned a value, *m*, which is defined as it’s position within this proximal-to-distal ordering, beginning with 0. Each gene is assigned a value, *n*, which is defined as the number of distinct APA sites that it contains. The expression (TPM) of every terminal fragment belonging to a given APA site is then summed to define the expression level of the APA site, and this process is repeated for every APA site within a gene. The expression level of each APA site is then scaled according to the following formula:

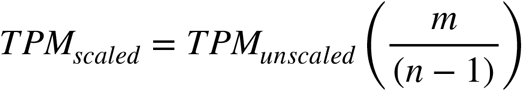

To quantify a gene’s relative APA site usage, we defined a term, *ψ*. Scaled and unscaled TPM values are summed across all APA sites within a gene, and ψ is defined as the ratio between these summed values:

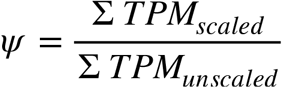

With this strategy, genes that show exclusive usage of the most gene-proximal APA site will be assigned a *ψ* value of 0, while those that show exclusive usage of the most gene-distal APA site will be assigned a *ψ* value of 1 (**Figure 1B**). Usage of both sites will result in a *ψ* value between 0 and 1 depending on the relative usage of the sites. Importantly, this strategy also applies to genes with more than 2 APA sites. In these cases, one *ψ* value is assigned to the entire gene without the need to do multiple pairwise comparisons between APA sites.

After calculating *ψ* values for genes in all samples, LABRAT compares *ψ* values of experimental replicates across experimental conditions to identify genes with statistically significantly different *ψ* values between conditions. This is done using a mixed linear effects model that tests the relationship between *ψ* values and experimental condition. A null model is also created in which the term denoting the experimental condition has been removed. A likelihood ratio test compares the goodness of fit of these two models to the observed data and assigns a p value for the probability that the real model is a better fit than the null model. In simple comparisons between two conditions, this approach mimics a t-test. However, this technique has the advantage of being able to easily incorporate covariates into significance testing. After performing this test on all genes, the raw p values are corrected for multiple hypothesis testing using a Benjamini-Hochsberg correction (Benjamini and Hochberg, 1995).

In addition, LABRAT determines whether a gene’s APA sites conform to either the tandem UTR or ALE structures (**Figure 1B**) and designates the gene accordingly. For genes with more than 2 APA sites, it is possible to contain both tandem UTR and ALE structures. These genes are designated as having a “mixed” APA structure.

### Identifying tissue-specific differences in APA with LABRAT

To demonstrate the ability of LABRAT to identify and quantify differences in APA, we analyzed RNAseq data from mouse brain and liver tissues (Li et al., 2017). Because neuronal tissues are known to be highly enriched for the use of distal APA sites (Miura et al., 2013), we reasoned that comparison of these two tissues might provide a positive control for LABRAT’s ability to identify differential APA.

We found 470 genes that displayed differential APA site usage between the tissues (FDR < 0.05) (**Figure 1C**). As expected, 68% of these genes showed increased usage of distal APA sites in brain, indicating a significant enrichment for the use of downstream APA sites in this tissue (binomial p = 3.2e-15). To further explore changes in *ψ* value for specific genes, we plotted read coverages over two genes that showed significantly more downstream APA site usage in brain tissue: Slc16a7 and Elavl1 (**Figure 1D, E**). For both genes, we observed significantly lower read coverages corresponding to usage of the distal APA site in the liver samples relative to the brain samples. Accordingly, LABRAT assigned these genes to have low *ψ* values in the liver samples, and high *ψ* values in the brain samples, indicating that LABRAT can accurately quantify APA.

To perform similar analyses in human samples, we analyzed over 5000 RNAseq samples from over 30 different human tissues produced as part of the Genotype-Tissue Expression (GTEx) project (GTEx Consortium, 2013). We quantified APA in these samples and observed relationships between tissue APA using PCA analysis (**Figure S1A**). In this analysis, brain and testis samples were clear outliers. Interestingly, performing the PCA analysis using only tandem UTR (**Figure S1B**) or ALE (**Figure S1C**) genes produced very similar results, suggesting that these two forms of APA are broadly coregulated across many tissues.

To understand more about APA in human brain and testis, we compared their APA profiles to those observed in human liver samples. As expected, we observed that brain samples exhibited a significant bias for the use of downstream APA sites (p < 2.2e-16) (**Figure S1D**). Conversely, testis samples exhibited a similar bias for the use of upstream APA sites (p < 2.2e-16) (**Figure S1E**). The propensity of testis to use upstream APA sites has been previously observed (Liu et al., 2007; Wang et al., 2008; Zhang et al., 2005) and is likely a key feature of spermatogenesis (Li et al., 2016). Overall, these results demonstrate the ability of LABRAT to recapitulate previously reported observations and gave us confidence in its results moving forward.

### Comparison of LABRAT to similar methods of APA quantification

To compare LABRAT with other APA analysis tools, we generated a synthetic RNAseq dataset containing 50 million reads in which 1250 genes displayed increased distal APA site usage, 1250 genes displayed increased proximal APA site usage, and 2500 genes displayed no change in APA site usage (Frazee et al., 2015). We used the software packages QAPA (Ha et al., 2018), DaPars (Xia et al., 2014), and Roar (Grassi et al., 2016) in addition to LABRAT to quantify APA in these data.

QAPA, like LABRAT, uses lightweight alignments to quantify APA. Reassuringly, we found that *ψ* values calculated by LABRAT were highly correlated to the analogous metric used by QAPA, PPAU (R = 0.81) (**Figure S1F**). In comparing the four methods, LABRAT was the best suited to accurately identify differential APA in the simulated data (**Figure 1F**). We further found that the accuracy of LABRAT was not noticeably affected by read depth down to one million reads (**Figure S1G**).

### Alternative polyadenylation isoforms are differentially localized in cell bodies and projections

Multiple studies have found that alternative polyadenylation decisions made during nuclear processing can influence the subcellular localization of the resulting transcript, particularly in neuronal cells (Ciolli Mattioli et al., 2019; Taliaferro et al., 2016; Tushev et al., 2018). However, it has been unclear how widespread this effect is and whether it was driven primarily by tandem UTRs or ALEs. To address this, we used LABRAT to analyze the relative APA status of 26 paired transcriptomic datasets from cell body and projection samples from neuronal cells, NIH 3T3 cells, and MDA-MB231 cells (Farris et al., 2019; Goering et al., 2019; Hudish et al., 2020; Mardakheh et al., 2015; Minis et al., 2013; Taliaferro et al., 2016; Tushev et al., 2018; Wang et al., 2017; Zappulo et al., 2017).

For all samples, we identified genes whose *ψ* value was significantly different between subcellular compartments (FDR < 0.05), finding between 10 and 740 genes that fit this criterion in each sample (**Figure 2A**). For these genes, we then compared their *ψ* values across compartments by subtracting the *ψ* value in the cell body from the *ψ* in the projection to define *Δψ*. Genes with positive *Δψ* values therefore had their distal APA isoform enriched in projections while those with negative *Δψ* values had their proximal APA isoform enriched in projections.

**Figure 2.**
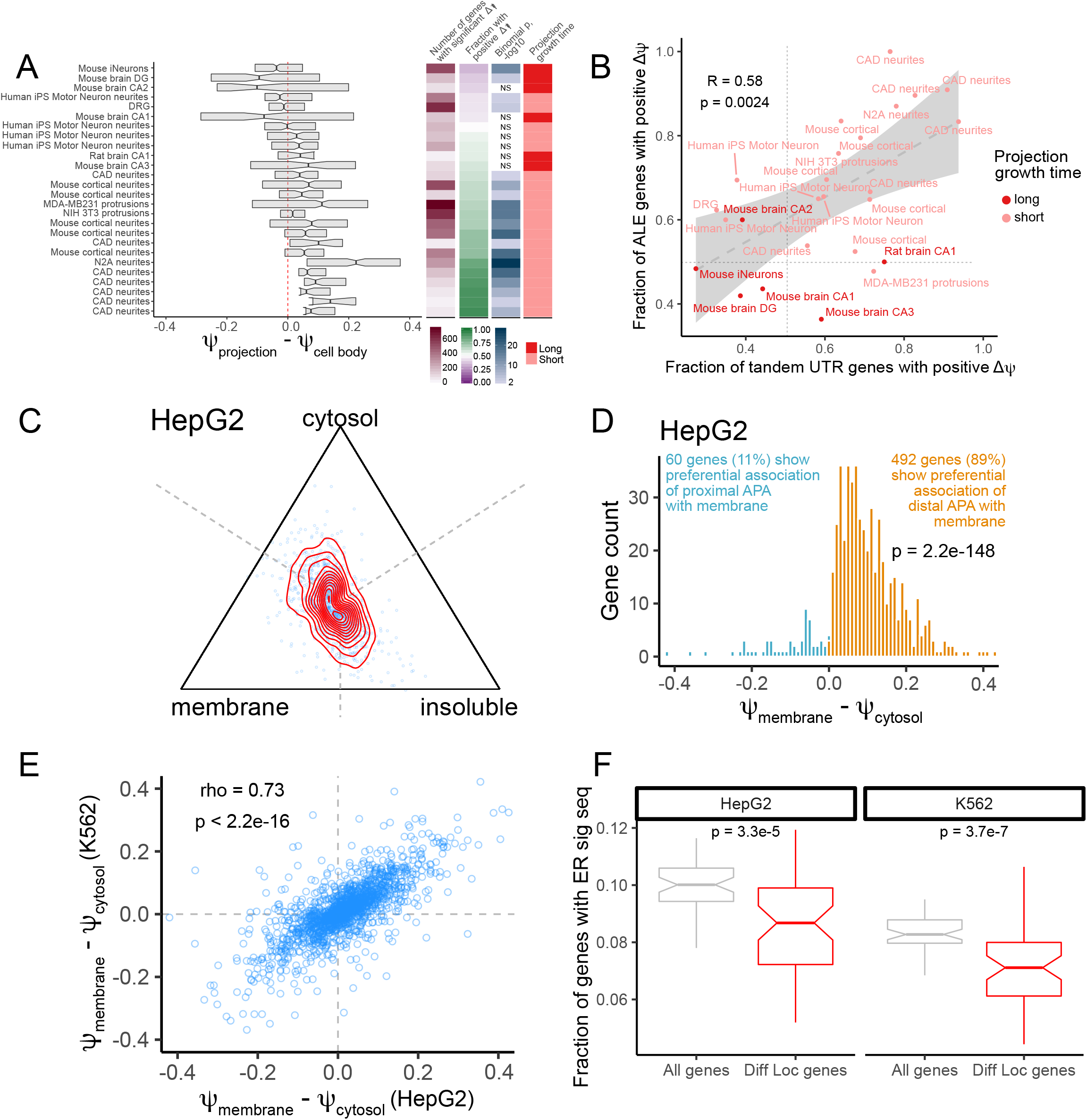
Alternative polyadenylation is associated with RNA localization in a variety of cell types. (A) Comparison of *ψ* values for RNA isolated from cell projections and cell bodies. *ψ* values for all genes were calculated using RNA collected from cell projection and cell body compartments, and genes with significantly different *ψ* values across compartments were identified (FDR < 0.05). *Δψ* values (cell projection – cell body) for these genes are indicated by boxplots. P values in blue represent binomial p values for deviations from the expected 50% chance for a gene to have a positive *Δψ* value. Samples were also separated according to the amount of time that projections were allowed to grow before their RNA content was analyzed. This is represented by the long (at least 6 days) and short (2 days or less) categories colored in red. (B) As in A, *ψ* values for all genes were calculated using RNA collected from cell projection and cell body compartments, and genes with significantly different *ψ* values across compartments were identified (FDR < 0.05). The fraction of significant tandem UTR and ALE genes with positive *Δψ* values were plotted on the x and y axes, respectively. (C) Simplex plot indicating *ψ* values calculated from RNA isolated from biochemically defined cytosolic, membrane-associated, and insoluble fractions of HepG2 cells. Genes with equal *ψ* values in all three fractions are represented by dots equidistant from each vertex (at the intersection of the dotted lines). Genes that displayed higher *ψ* values in a given fraction than the others are represented by dots placed closer to that fraction’s vertex. Red lines indicate the density of dots. (D) Comparison of *ψ* values in HepG2 cytosolic and membrane fractions for genes whose *ψ* value was significantly different between these compartments (FDR < 0.01). (E) Correlation of *Δψ* values (membrane – cytosol) for all genes expressed in both HepG2 and K562 cells. (F) Fraction of genes with nonsignificant *Δψ* values (membrane vs. cytosol, gray) and those with significant *Δψ* values (red) that encode peptides that have ER signal sequences as defined by SignalP. Distributions of this fraction were created through bootstrapping in which 40% of the genes were sampled 100 times. P values were calculated using a Wilcoxon rank sum test.

We found that for 19 of these 26 samples, over 50% of significant genes had positive *Δψ* values, indicating a broad connection between the use of distal APA sites and localization of the resulting transcript to cell projections (**Figure 2A**). Further, we observed a relationship between the amount of time that the projection had been allowed to grow and the fraction of genes with positive *Δψ* values. Of the samples in which the projections had grown for 2 days or less, 15 out of 20 showed a significant bias for the association of distal APA sites with projections. Conversely, of the samples in which the projections had grown for 6 days or more, 0 out of 6 showed a significant bias for the association of distal APA sites with projections. This suggests that distal APA transcripts may play a role in early projection outgrowth but may be less important in mature projections.

Given the conflicting reports about the relative contributions of distal APA produced by tandem UTR and ALEs to the transcriptomes of cell projections (Ciolli Mattioli et al., 2019; Taliaferro et al., 2016; Tushev et al., 2018), we analyzed these two classes of APA isoforms separately. Across the 26 subcellular comparisons, we found a strong, significant correlation (R = 0.58, p = 0.0024) between the fraction of ALE genes with positive *Δψ* values and the fraction of tandem UTR genes with positive *Δψ* values (**Figure 2B**). This indicates that both classes of genes are preferentially contributing their distal APA isoforms to projections and suggests that these two classes of alternative poly(A) site selection may be regulated by a common mechanism.

### Alternative polyadenylation isoforms are differentially localized in biochemically defined cytosolic and membrane fractions

To further explore connections between APA and RNA localization beyond cell projections, we used LABRAT to analyze RNAseq data from a biochemical fractionation of 3 cell types, Drosophila DM-D17-C3 (D17) cells, human HepG2 cells, and human K562 cells (Benoit Bouvrette et al., 2018). In these data, cells were fractionated into nuclear, cytosolic, membrane-associated and insoluble fractions. RNA was isolated from each of these fractions and prepared for high-throughput sequencing using either polyA-selection-based or ribosomal RNA-depletion-based library preparation. For each fraction, two replicates of each library preparation method were sequenced.

As with the projection data, we compared *ψ* values for genes across cellular compartments. Hierarchical clustering of samples based on *ψ* values revealed that samples from the same fraction generally clustered with each other, indicating the high quality of the data. indicating the quality of the data (**Figure S2A-C**). To minimize the effect of library preparation on the identification of genes with significantly different *ψ* values across compartments, we included the library preparation method as a covariate in LABRAT’s linear model. This allowed us to pool all of the samples for a given compartment in order to identify genes with significantly different *ψ* values between compartments regardless of library preparation method.

We first identified genes with significantly different *ψ* values across any pairwise comparison between cytosolic, membrane-associated, and insoluble fractions (FDR < 0.05). Based on our observations relating distal APA and RNA localization to projections, we then asked if any of these fractions were associated with higher *ψ* values than the other two. We visualized these comparisons using simplex plots (**Figure 2C**). In these plots, each dot represents a gene, and its position is determined by the relative *ψ* values in each fraction. A gene with a *ψ* value of 1 in a fraction and *ψ* values of 0 in the other two would be placed at that fraction’s vertex while a gene with equal *ψ* values in all 3 fractions would be placed equidistant from each vertex at the intersection of the dotted lines. We found that genes tended to have higher *ψ* values in the membrane fraction (**Figure 2C, S2D, E**), indicating a preferential association of downstream APA isoforms with that fraction.

Because of this observation, we then focused on comparing the cytosolic and membrane fractions. When comparing the cytosolic and membrane fractions of HepG2 cells, we identified 552 genes that had significantly different *ψ* values between the fractions (FDR < 0.01). Of these, 492 (89%) had a *ψ* value that was higher in the membrane fraction than the cytosolic fraction, indicating a broad association between transcripts produced using distal APA sites and the membrane fraction (**Figure 2D**). We observed highly similar results when comparing the cytosolic and membrane fractions from K562 cells and D17 cells (**Figure S2F, G**).

We then queried whether the same genes had differential APA isoform associations with the cytosolic and membrane fractions in the HepG2 and K562 samples. To test this, we calculated *Δψ* values (membrane – cytosol) for all genes expressed in both cell lines. We observed a strong correlation (R = 0.73) between *Δψ* values in the two cell lines (**Figure 2E**), suggesting that the effects of APA on transcript membrane association are shared between cell lines and are therefore likely transcript-specific with a conserved mechanistic basis.

The ER comprises a large fraction of cellular membranes, and RNA localization to the ER is important for cotranslational access to the secretory pathway. We therefore asked whether transcripts with significant membrane vs. cytosol *Δψ* values were more or less likely than expected to encode the peptide-based signal sequences required for RNA transport to the ER through cotranslational targeting. We identified ER signal sequences using SignalP (Almagro Armenteros et al., 2019). Interestingly, we found that in both the HepG2 and K562 samples, genes that had significant membrane vs. cytosol *Δψ* values were significantly less likely to contain an ER signal sequence than other genes (**Figure 2F**). This observation therefore suggests two alternative modes of RNA localization to the ER: one for transcripts that encode signal peptides and another for those that do not. Specifically, mRNAs that are not cotranslationally targeted by signal peptide recognition appear to be targeted by a mechanism involving distal APA use.

### The transcription speed of RNA polymerase II regulates alternative polyadenylation site choice

The speed of transcription by RNA Polymerase II (Pol II) regulates multiple co-transcriptional processes, including alternative splicing, and termination that is coupled to poly(A) site cleavage (Cortazar et al., 2019; Dujardin et al., 2014; Fong et al., 2014; Jonkers et al., 2014; de la Mata et al., 2003). To assess how changes in Pol II speed can affect APA, we used LABRAT to analyze RNAseq samples from HEK293 cells that expressed either wildtype or slow Pol II (Fong et al., 2014). The slow Pol II mutant used in these studies is a single amino acid substitution in the funnel domain of the Pol II large subunit Rpb1 (R749H).

During transcription, a gene-proximal APA site is necessarily transcribed before a gene-distal APA site. There exists a time, therefore, during which the proximal site is the only APA site that exists on the transcript. Reducing the speed of Pol II transcription would increase this time in which the proximal site is free from competition with the distal site. We hypothesized that this would lead to an increase in usage of the proximal APA site (**Figure 3A**). Indeed, we found that for many genes, proximal APA site usage was increased in slow Pol II samples (**Figure 3B**), and that overall there was a shift towards increased usage of the proximal site (**Figure 3C**).

**Figure 3.**
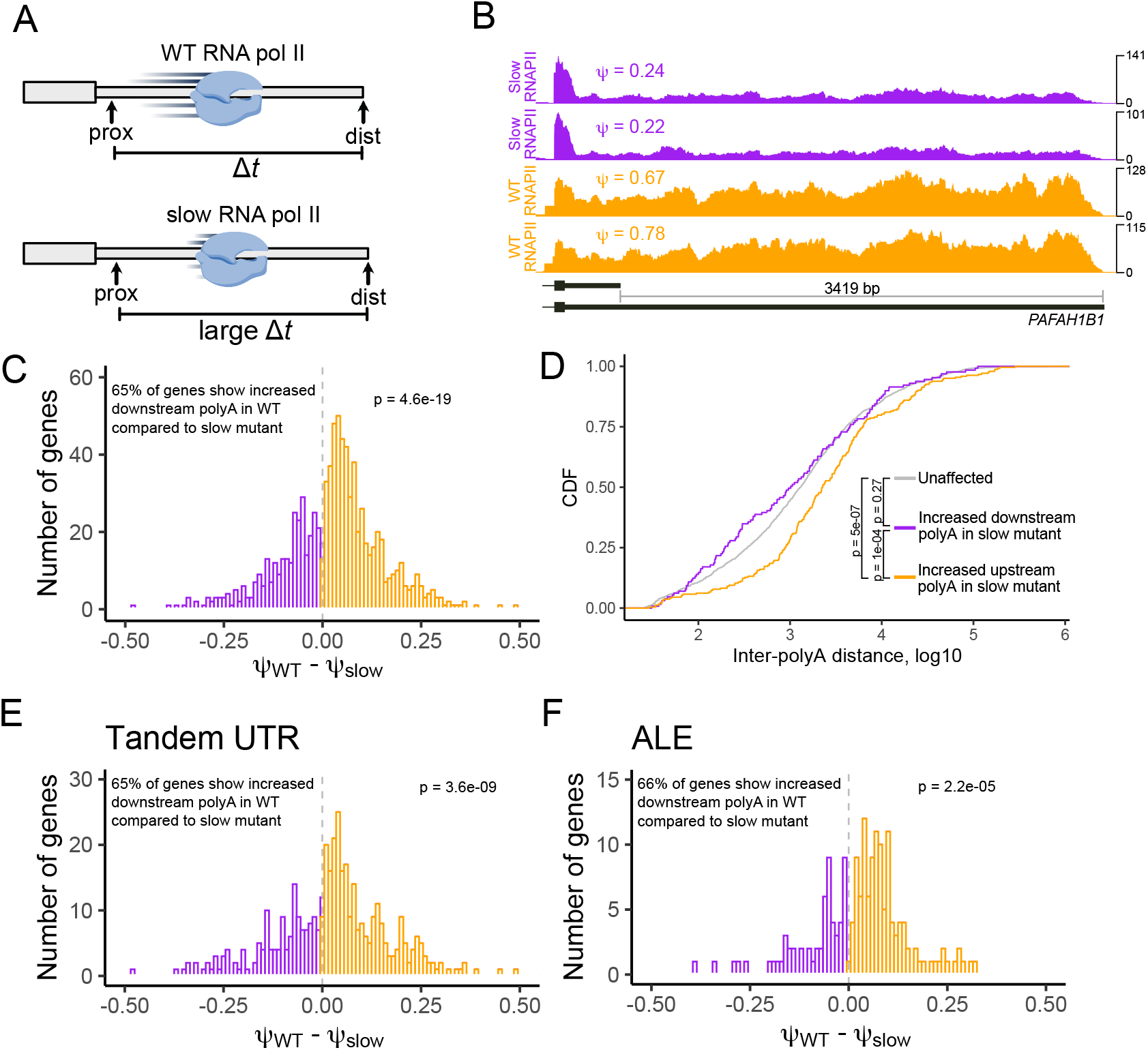
The speed of RNA polymerase II influences APA. (A) Model for how polymerase speed can affect alternative polyadenylation. During the time between transcription of proximal and distal polyadenylation sites, the proximal site can be recognized and used but the proximal site cannot. Increasing this time of proximal site exclusivity by decreasing the speed of RNA polymerase may increase the likelihood of the proximal site being used. (B) Read coverage and *ψ* values for the gene *PAFAH1B1* in cells expressing wildtype (orange) and slow (purple) RNA polymerase II. (C) Comparison of *ψ* values in cells expressing wildtype and slow RNA polymerase II for genes whose *ψ* value was significantly different between these samples (FDR < 0.05). (D) Distance between alternative polyadenylation sites for genes that displayed increased upstream APA (orange), increased downstream APA (purple), or whose APA did not change (gray) in cells expressing a slow RNA polymerase II compared to cells expressing wildtype RNA polymerase II. (E-F) As in D, comparison of *ψ* values in cells expressing wildtype and slow RNA polymerase II for tandem UTR (E) and ALE (F) genes whose *ψ* value was significantly different between these samples (FDR < 0.05).

If the shift in APA was related to the amount of time during which the proximal site was exclusive, then the shift should be most pronounced in genes in which the distance between proximal and distal sites is large. Consistent with this hypothesis, we found that this “inter-polyA distance” for genes that displayed increased proximal APA was significantly longer than expected (**Figure 3D**), further suggesting that changes in Pol II kinetics can predictably alter APA.

If alternative polyadenylation of tandem UTRs and ALEs were generally coregulated, then it would be expected that changes in Pol II speed would affect both classes of genes. To test this, we examined the increase in proximal APA site usage caused by slow transcription in the context of tandem UTR and ALE genes separately. We found that proximal APA usage was increased for both tandem UTR and ALE genes (**Figure 3E, F**), indicating that the two classes of genes are similarly affected by changes in Pol II speed and consistent with the idea that they are coregulated by a common mechanism.

### Dozens of RNA-binding proteins (RBPs) regulate relative APA isoform abundance across many genes

To investigate the contributions that individual RBPs can have to the regulation of APA isoform abundance, we analyzed the ENCODE RBP knockdown RNAseq datasets with LABRAT (Consortium, ENCODE Project et al., 2012; Davis et al., 2018). This resource contains 523 shRNA-mediated RBP knockdown RNAseq experiments spread across human HepG2 and K562 cell lines. We compared *ψ* values for all expressed genes between RBP knockdown and control knockdown samples for 191 RBPs that were expressed in both cell lines. To identify genes that had significantly different *ψ* values (FDR < 0.05) between RBP knockdown and control knockdown samples, we incorporated the cell line of the experiment as a covariate in LABRAT’s linear model.

We began by assessing the reproducibility of changes in APA isoform abundance upon RBP knockdown between the two cell lines. To do this, we correlated *Δψ* values (control knockdown – RBP knockdown) for all expressed genes in a given RBP knockdown in HepG2 cells with their *Δψ* values upon knockdown of the same RBP in K562 cells. We therefore end up with one correlation coefficient per RBP knockdown. As a control, we compared these values to correlations of *Δψ* values where the RBP that was knocked down was different between the cell lines (**Figure 4A**). Reassuringly, we found that correlations between experiments in which the expression of the same RBP was knockdown were significantly higher than those in which the expression different RBPs were knocked down (p = 1.5e-19, Wilcoxon ranksum test). When we restricted the comparison to genes that had significantly different *ψ* values between RBP and control knockdowns (FDR < 0.05), we observed a much higher correlation of *Δψ* values between cell lines (**Figure 4A**). These results gave us confidence that we could accurately quantify APA isoform abundance in the ENCODE datasets.

**Figure 4.**
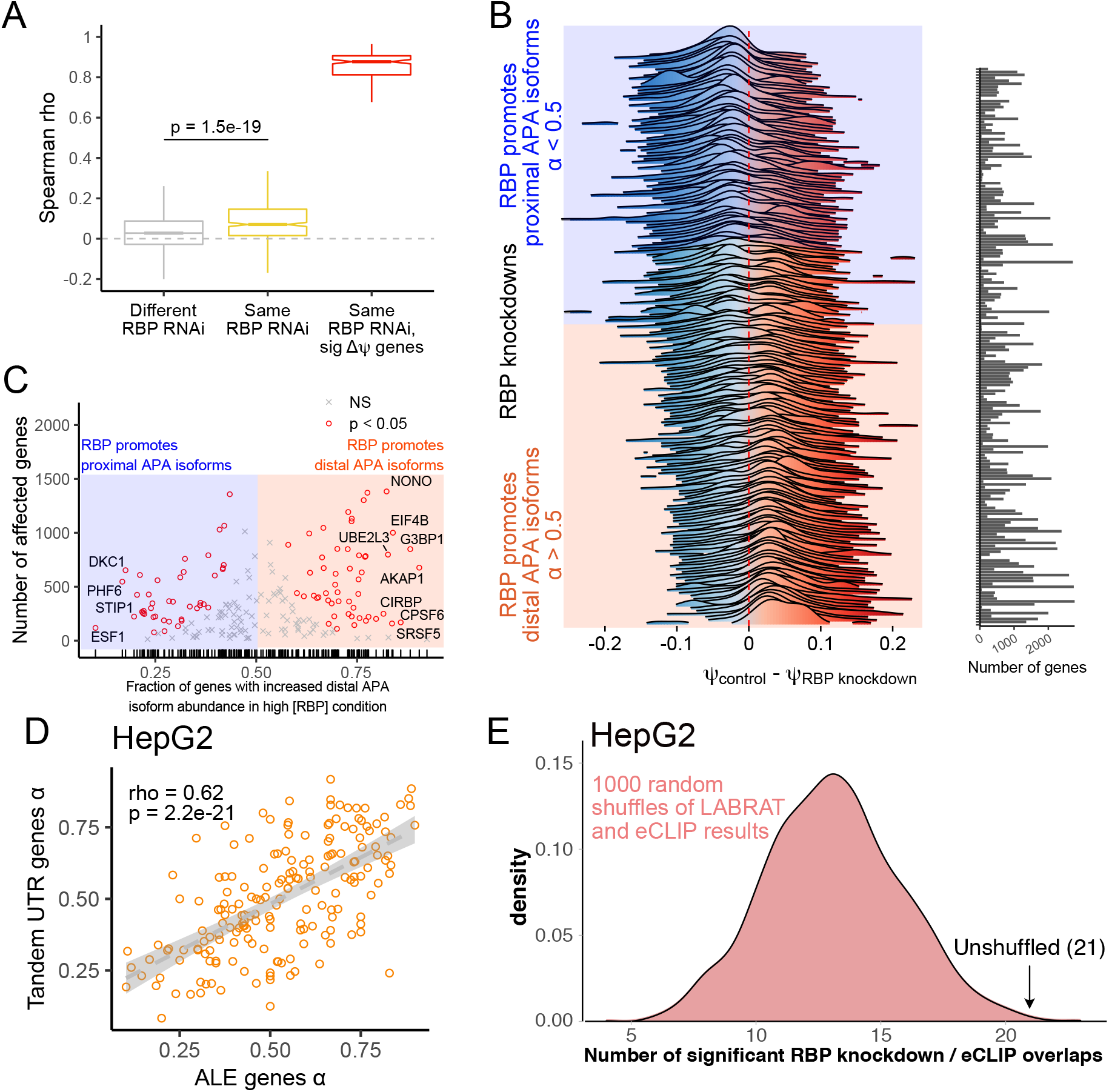
Many RBPs promote proximal or distal APA isoform abundance in hundreds of genes. (A) Correlation of all *ψ* values across HepG2 and K562 cell lines for all ENCODE RBP-knockdown RNAseq experiments. In gray, correlation coefficients for comparisons of different RBP knockdowns are shown (e.g. RBP X in HepG2 vs. RBP Y in K562). In yellow, correlation coefficients for comparisons of the same RBP knockdown are shown (e.g. RBP X in HepG2 vs RBP X in K562). In red, this comparison is restricted to only those genes whose *ψ* value significantly differed between the RBP knockdown and control knockdown samples (e.g. RBP X in HepG2 vs RBP X in K562, significant *Δψ* genes only). In identification of these significant genes, the cell line was included as a covariate. (B) Comparison of *ψ* values in RBP knockdown and control samples for genes whose *ψ* value was significantly different between these samples (FDR < 0.05). The number of genes with significant *Δψ* values in each comparison is indicated by the bar graph. A term, *α*, was defined as the fraction of these genes that displayed higher *ψ* values in the high RBP state (control knockdown) versus the low RBP state (RBP knockdown). (C) For each RBP knockdown, the number of genes with significant *Δψ* values (FDR < 0.05) is indicated on the y axis while the fraction of these genes with positive *Δψ* values (control knockdown – RBP knockdown) is indicated on the x axis. Knockdowns whose fraction of genes with positive *Δψ* values significantly differs from the expected 50% are indicated with red circles. (D) *α* values for each RBP knockdown in HepG2 cells were calculated using tandem UTR and ALE genes independently. These were then plotted and correlated. Each dot in this plot represents one RBP knockdown experiment. (E) Among 84 RBPs expressed in HepG2 cells, overlaps between the genes whose APA was sensitive to RBP knockdown and the genes whose 3′ UTRs were bound by the RBP in eCLIP experiments were calculated. The significance of this overlap was calculated using a binomial test. 21 RBPs bound the 3′ UTRs of their APA targets more often than expected (binomial p < 0.05). To assess whether this was more than the expected number of significant RBPs, relationships between RBPs and their lists of APA and eCLIP targets were shuffled 1000 times, and the analysis was repeated after each shuffle to create a null distribution (pink).

For each RBP knockdown experiment, we then took the genes with significantly different *ψ* values between RBP and control knockdowns and analyzed the distribution of their *Δψ* values (control knockdown – RBP knockdown) (**Figure 4B**). We observed that many RBP had distributions of *Δψ* values that were skewed towards being mostly positive or mostly negative. We defined a term, *α*, as the fraction of these genes with positive *Δψ* values. RBPs with *α* values greater than 0.5 therefore were broadly associated with increased distal APA isoform abundance while those with *α* values less than 0.5 were associated with increased proximal APA isoform abundance. 94 RBPs had *α* values that were significantly skewed from the expected value of 0.5 (binomial p < 0.01), and of these 52 had *α* values of greater than 0.5 while 42 had *α* values less than 0.5 (**Figure 4C**).

For each RBP knockdown experiment we then calculated *α* values for tandem UTR and ALE genes separately. *α* values for these two APA types were highly correlated (R = 0.62), further indicating that these two mechanisms of APA regulation are not independent and share elements in common (**Figure 4D, Figure S3A**).

If changes in APA isoform abundance upon RNAi were directly due to loss of the RBP, then we would expect that the RBP would directly bind the 3′ UTRs of the genes whose APA it regulates. To test this, we analyzed RBP/RNA interactions as measured by the eCLIP experiments performed as part of the ENCODE project (Van Nostrand et al., 2016). We observed that some RBPs displayed highly promiscuous 3′ UTR binding while others bound very few 3′ UTRs (**Figures S3B, C**).

In HepG2 cells, 84 RBPs had both RNAseq data from RNAi experiments and eCLIP data. For each RBP, we calculated how many of the genes with significant changes in *ψ* value upon RBP knockdown also contained an eCLIP peak for that RBP in their 3′ UTR. We then calculated whether this overlap of RBP binding and function was statistically significant (binomial p < 0.05). For 21 of these RBPs, we observed a significant overlap between the RBPs functional APA targets and the 3′ UTRs it bound (**Figure 4E**). To assess whether this was more or less than the number of expected significant RBPs, we shuffled the relationships between RBPs and their lists of APA targets and bound 3′ UTRs and again calculated the number of RBPs that showed significant overlap between APA and eCLIP data. Repeating this process 1000 times gave us a null distribution of the expected number of RBPs with significant overlaps and indicated that the observed number of overlaps was significant in HepG2 cells (p = 0.006).

Although we did not observe a similar significant relationship between APA and eCLIP data in K562 cells (p = 0.4) (**Figure S3D**), overall, these results indicate that many of the RBPs tested are modulating relative APA isoform abundance through direct interactions.

### Misregulation of alternative polyadenylation is cancer type specific and correlates with patient survival

Changes APA have long been known to be associated with cancer (Masamha and Wagner, 2018; Yuan et al., 2019). Most often, APA is thought to contribute to cancer phenotypes through a general increased usage of proximal APA sites, which are thought to be associated with increased expression of oncogenes and proliferation of cell lines (Mayr and Bartel, 2009; Sandberg et al., 2008). To further explore this phenomenon, we used LABRAT and data from The Cancer Genome Atlas (TCGA) (Cancer Genome Atlas Research Network et al., 2013) to examine changes in APA between matched tumor and normal samples from 671 patients across 21 different cancers.

For each cancer, we identified between 130 and 3043 genes that displayed significant differences in *ψ* values (FDR < 0.05) between tumor and normal samples. We then defined *Δψ* values (tumor – normal) to ask whether proximal or distal sites showed increased usage in tumor samples. For some cancers, including Lung Squamous Cell Carcinoma (LUSC), Urothelial Bladder Carcinoma (BLCA) and Lung Adenocarcinoma (LUAD), tumors displayed the expected pattern of increased proximal APA in tumors (**Figure 5A**). Conversely, Thyroid Cancer (THCA) and Kidney Renal Clear Cell Carcinoma (KIRC) showed strong biases in the opposite direction with increased distal APA in tumors. Mechanisms that drive APA dysregulation are therefore likely specific to different cancer types, and it is not true that increased proximal APA is a general feature of cancer cells.

**Figure 5.**
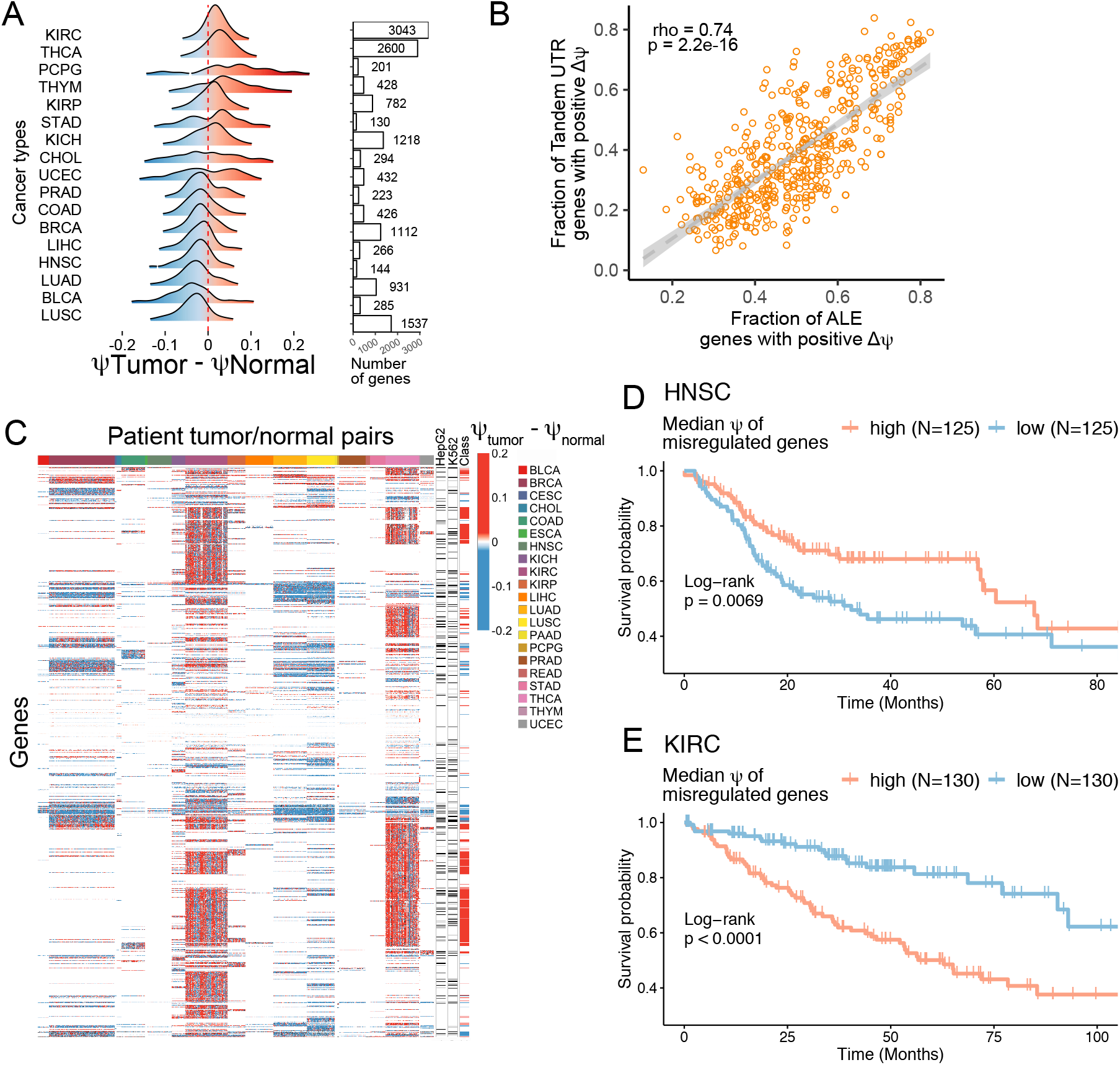
Misregulation of alternative polyadenylation in primary tumors. (A) Comparison of ψ values in matched patient tumor and control samples for genes whose ψ value was significantly different between these samples (FDR < 0.05). The number of genes with significant *Δψ* values in each comparison is indicated by the bar graph. (B) As in A, *ψ* values for all genes were calculated in matched patient tumor and normal tissue samples, and genes with significantly different *ψ* values across samples within a cancer type were identified (FDR < 0.05). The fraction of significant tandem UTR and ALE genes with positive *Δ-ψ* values were plotted on the x and y axes, respectively. Each dot represents one patient sample pair. (C) Genes with significantly different ψ values across samples within a cancer type (FDR < 0.05) are colored according to their *Δψ* value (tumor – control). Columns represent matched patient samples while rows represent genes. Black ticks (right) represent whether or not the gene displayed a significantly different *ψ* value (FDR < 0.05) between biochemically defined cytosolic and membrane-associated fractions in HepG2 and K562 cells (Figure 2D). Genes were further separated into classes of those with increased *ψ* values in KIRC and THCA tumor samples (red ticks, right) and those with decreased *ψ* values in BRCA, LUAD and LUSC tumor samples (blue ticks, right). (D-E) Survival analysis for APA misregulation in head and neck squamous cell carcinoma and kidney renal clear cell carcinoma, respectively. Patients were grouped into extreme quartiles by ranked median *ψ* values for misregulated genes as defined in Figure 5A for the respective tumors. In figure 5A, HNSC tumors were associated with decreased *ψ* values. Here, lower *ψ* values are associated with poor prognosis. Conversely, in figure 5A KIRC tumors were associated with increased *ψ* values, and here, increased *ψ* values are associated with poor prognosis.

We then compared *ψ* values in the TCGA data for tandem UTR genes and ALE genes separately. For each pair of tumor and normal samples, we calculated the fraction of genes with significantly different *ψ* values across conditions (FDR < 0.05) in which the *ψ* value was greater in the tumor sample than the normal sample. Put another way, for each patient, we calculated the fraction of significant tandem UTR and ALE genes with positive *Δψ* (tumor – normal) values (**Figure 5B**). The tandem UTR- and ALE-derived fractions were strongly correlated with each other (R = 0.74), again suggesting that these two modes of APA may be coregulated.

We wondered if APA was misregulated in the same genes across many different cancer types or whether the set of genes with misregulated APA was cancer type specific. Although many APA misregulated genes were specific to certain cancers, we did observe that hundreds of genes repeatedly showed misregulation across multiple cancers (**Figure 5C**). We defined a set of genes that repeatedly showed increased proximal APA usage in BLCA, LUAD, and LUSC tumors. Using gene ontology analysis, we found that these genes were significantly enriched for those encoding singlestranded RNA binding proteins (Eden et al., 2009). Similarly, we defined a set of genes that repeatedly showed increased distal APA usage in THCA and KIRC. These genes were enriched for being involved in programmed cell death and responses to stress.

We enquired whether transcripts we identified whose APA status correlates with membrane association (**Figure 2C, D**) are among those subject to misregulation in tumors. Many of these membrane associated mRNAs showed significantly different *ψ* values between tumor and normal samples, suggesting that the subcellular localization of these transcripts may be altered in cancerous cells.

To determine if the degree of APA misregulation was related to patient prognosis, we performed survival analyses for patients from the TCGA dataset. In figure 5A, we defined genes with tumorspecific APA misregulation by comparing *ψ* values in tumor and matched normal patient samples. For each tumor, we then calculated a median *ψ* value across these genes in thousands of tumor RNAseq samples. Using this median *ψ* of misregulated genes, we ranked patients and separated them into quartiles. The extreme quartiles (patients with the highest and lowest *ψ* values for misregulated genes) for each cancer were compared. We found that for head and neck squamous cell carcinoma (HNSC), a cancer that typically exhibits increased proximal APA, patients with lower *ψ* values in misregulated genes had poorer prognoses (p = 0.0069) compared to patients with higher *ψ* values for the same genes (**Figure 5D**). Conversely, for kidney renal clear cell carcinoma (KIRC), a cancer that typically exhibits increased distal APA, we found the opposite. Patients with lower *ψ* values in misregulated genes had better outcomes compared to patients with higher *ψ* values (p < 0.0001) (**Figure 5E**). Therefore, the direction of APA misregulation is cancer-specific, and both increased proximal and distal APA are associated with poor patient prognosis, depending on the cancer type.

### Usage of distal APA sites is broadly but weakly associated with decreased RNA expression

Some of the original studies on the relationship between APA and RNA expression reported that distal APA is associated with a decrease in RNA levels (Mayr and Bartel, 2009) while more recent genome-wide studies have reported that the relationship is less clear (Spies et al., 2013; Venkat et al., 2020). To comprehensively examine the relationship between APA and gene expression, we compared changes in *ψ* and changes in RNA levels across the 191 ENCODE RBP knockdown sample pairs and the 671 TCGA tumor/normal sample pairs. To do so, we defined a term, rho (*ρ*), as the correlation between changes in *ψ* and changes in gene expression across two samples (**Figure 6A**). Sample comparisons where *Δψ* and gene expression changes are positively correlated indicate that distal APA and increased RNA levels were associated, and these comparisons will have positive *ρ* values. Conversely, sample comparisons where *Δψ* and gene expression changes are negatively correlated indicate that distal APA and decreased RNA levels were associated, and these comparisons will have negative *ρ* values.

**Figure 6.**
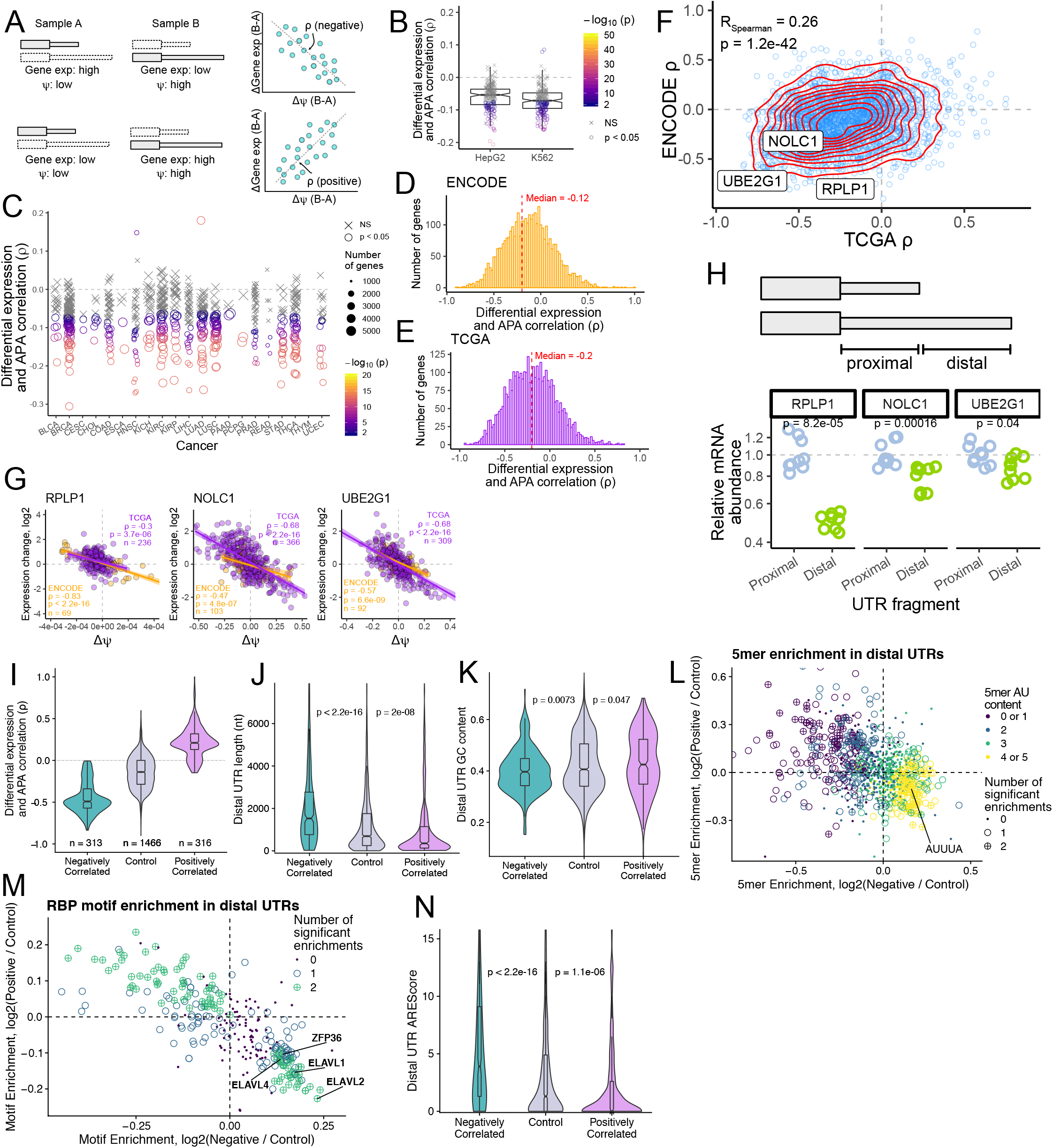
Comprehensive analyses of connections between alternative polyadenylation and transcript expression. (A) Diagram of correlation between APA and transcript expression. Rho (*ρ*) is defined as the correlation between changes in gene expression and changes in *ψ* value across two conditions. In the scenario described in the top row, the overall RNA expression level for the gene is high in sample A but low in sample B while the gene’s *ψ* value is low in sample A and high in sample B. Changes in gene expression and *Δψ* are therefore negatively correlated, giving *ρ* a negative value. Conversely, in the scenario described in the bottom row, changes in gene expression and *ψ* are positively correlated. (B) *ρ* values across all expressed genes within a comparison for the ENCODE RBP knockdown data. Each dot represents a single comparison (RBP knockdown vs control knockdown). P values for the correlation between gene expression and APA are indicated by dot shape and color. (C) *ρ* values across all expressed genes with a comparison for the TCGA paired tumor/control sample data. Each dot represents a single patient’s tumor and control samples. P values for the correlation between gene expression and APA are indicated by dot shape and color. (D) Gene-level *ρ* values across all ENCODE RBP knockdown experiments. (E) Gene-level *ρ* values across all TCGA tumor/control sample pairs. (F) Correlation of gene-level *ρ* values derived from the ENCODE and TCGA datasets (D and E). Red lines indicate the density of points, and the locations of three genes selected for further study are indicated by labels. (G) Correlation between gene expression changes and *Δψ* for three genes. Orange dots represent ENCODE sample pairs (RBP knockdown vs. control knockdown) while purple dots represent TCGA sample pairs (tumor vs. control samples). (H) Top: illustration of the UTR fragments fused to the Firefly luciferase gene. Bottom: RT-qPCR-derived relative levels of firefly luciferase mRNA expression when the proximal and distal UTR fragments of the indicated genes were fused. Values indicate ratios between the abundances of Firefly and Renilla luciferase mRNAs with this ratio in the proximal UTR comparison set to 1. P values were calculated using a Wilcoxon ranksum test. (I) Correlation between gene expression changes and *Δψ* was used to define positively correlated, negatively correlated and control genes with two APA isoforms. Correlations are calculated for ENCODE and TCGA separately. (J) Distal UTR lengths of each gene set. P values were calculated using a Wilcoxon ranksum test. (K) Distal UTR GC content of each gene set. P values were calculated using a Wilcoxon ranksum test. (L) Five-mer enrichments in the distal 3′ UTRs of positively and negatively correlated gene sets vs control. Five-mers are significantly enriched (BH-adjusted p<0.05, Fisher’s exact test) in either both comparisons, one comparison or neither and are represented by a circle plus, open circle or closed dot respectively. Five-mers are colored by their AU content as ranked 0-5. Canonical AU rich element (ARE) “AUUUA” is highlighted as enriched in negatively correlated distal UTRs. (M) RBP motif enrichments in the distal 3′ UTRs of positively and negatively correlated gene sets vs control. RBP motifs are significantly enriched (BH-adjusted p<0.05, Fisher’s exact test) in either both comparisons, one comparison or neither and are represented by a green circle plus, blue open circle or purple dot respectively. Canonical ARE binding protein motifs are highlighted as enriched in negatively correlated distal UTRs. (N) Distal UTR AREScores of each gene set as calculated by AREScore software. P values were calculated using a Wilcoxon ranksum test.

We calculated *ρ* values across all genes for each RBP knockdown in the ENCODE data. In both the HepG2 and K562 samples, these *ρ* values overwhelmingly tended to be negative, but weakly so (**Figure 6B**). We similarly calculated *ρ* values across all genes for every patient-derived tumor/ normal pair in the TCGA data (**Figure 6C**). Again, these *ρ* values were consistently but weakly negative. These results indicate that although distal APA is generally associated with decreased gene expression, its contribution to changes in RNA levels is modest when comparing all genes in aggregate.

It could be the case, though, that for specific genes, APA and gene expression may be more strongly linked. To explore this, we calculated *ρ* values for each gene individually across all of the ENCODE and TCGA sample pairs (**Figure 6D, E**). The median genes again had weakly negative *ρ* values (−0.12 in the ENCODE data, −0.20 in the TCGA data). ENCODE- and TCGA-derived *ρ* values for genes were correlated with each other (**Figure 6F**). Tandem UTR genes and ALE genes displayed similar distributions of *ρ* values, indicating that relationships between gene expression and APA are of approximately equal strength in these two APA classes (**Figure S4A-D**).

The tails of the *ρ* value distributions were long, indicating that there were genes whose changes in *ψ* value and changes in expression were highly correlated across conditions. We selected three of these, RPLP1, NOLC1, and UBE2G1, for further analysis. Given that each of these genes had strong negative *ρ* values in both the ENCODE and TCGA data (**Figure 6G**), we reasoned that there may be elements in their distal UTRs downstream of the proximal APA site that confer reduced steady-state RNA levels. To test this experimentally, we fused the proximal and distal UTRs of each of these genes to the coding region of Firefly luciferase. Each construct was then site-specifically incorporated into the genome of HeLa cells through Cre-mediated recombination (Khandelia et al., 2011). The Firefly luciferase transcripts were coexpressed from a bidirectional tet-On promoter with unmodified Renilla luciferase. The RNA level of each Firefly-UTR fusion was measured using Taqman qRT-PCR with the Renilla luciferase transcript as a normalizing control. For all three tested genes, fusion of the distal UTR to Firefly luciferase significantly reduced the steady-state level of the RNA relative to a fusion with the proximal UTR, indicating that sequence elements downstream of the proximal APA sites likely have a role in reducing RNA expression (**Figure 6H**). We conclude that by comparing changes in gene expression and APA, we can identify functional elements within 3′ UTRs that regulate mRNA abundance.

### Features enriched in UTRs associated with gene expression changes

To better understand sequence elements downstream of proximal APA sites that may reduce RNA expression, we used the *ρ* values calculated for individual genes using ENCODE and TCGA sample sets to assign genes to positively correlated, negatively correlated or not correlated (control) gene sets (**Figure 6I, Figure S4E**). These gene sets behave differently: positively correlated genes are more highly expressed when downstream PAS are used (increased *ψ*) while negatively correlated genes become less highly expressed as they utilize more downstream PAS.

This analysis was simplified by only considering genes with two APA isoforms such that RNA expression could be explained by proximal or distal UTR usage. The analyzed UTR sequences were unique, meaning that tandem UTRs were separated into proximal and distal UTRs such that distal UTRs lacked their shared 5’ sequence (**Figure 6H**). This allowed us to identify sequence characteristics of distal UTRs that explain the differences in RNA expression of the positively correlated and negatively correlated gene sets.

Negatively correlated genes were found to have longer distal UTRs with lower GC content than expected (**Figure 6J, 6K**). Additionally, they were generally enriched for AU rich five-mers including the canonical AU rich element (ARE) “AUUUA” (**Figure 6L, Figure S4F**). Conversely the distal UTRs of positively correlated genes were depleted for AU-rich five-mers (**Figure S4G**). Unsurprisingly given their AU-richness, negatively correlated genes were enriched for ARE binding protein motifs in their distal UTRs and contained more AREs as scored by AREScore (Spasic et al., 2012) (**Figure 6M, 6N**). AREs are destabilizing RNA elements bound by several ARE binding proteins that facilitate RNA degradation. The presence of AREs in distal UTRs of negatively correlated genes is consistent with lower RNA expression when downstream PAS are utilized. It is important to note that the distal UTRs of positively correlated genes are depleted for AREs consistent with their higher expression. These results suggest that APA can regulate gene expression through the inclusion of destabilizing AREs in a transcript’s 3’UTR.

### Regulatory effects of RBPs on APA isoform abundance inferred from ENCODE data can be observed in TCGA data

The relation between RBP expression and the widespread misregulation of APA in cancer cells is poorly understood. We investigated this problem by examining expression in patient samples of the 191 RBPs that potentially influence APA isoform abundance revealed by our analysis of ENCODE knockdown RNAseq results (**Figure 4B, C**). Based on the ENCODE RBP knockdown data, we defined *α* values for RBPs where values of greater than 0.5 indicated an RBP that promoted distal APA isoform abundance while values of less than 0.5 indicated an RBP that promoted proximal APA isoform abundance. To compare *α* values to RBP effects on APA isoform abundance observed in the TCGA data, we defined another term, *β*, as the correlation between the change in RNA expression of an RBP between tumor and matched normal TCGA samples and the median *Δψ* of genes with significantly different APA between the samples (FDR < 0.05) (**Figure 7A**). RBPs with positive *β* values are therefore associated with increased distal APA isoform abundance while those with negative *β* values are associated with increased proximal APA isoform abundance.

**Figure 7.**
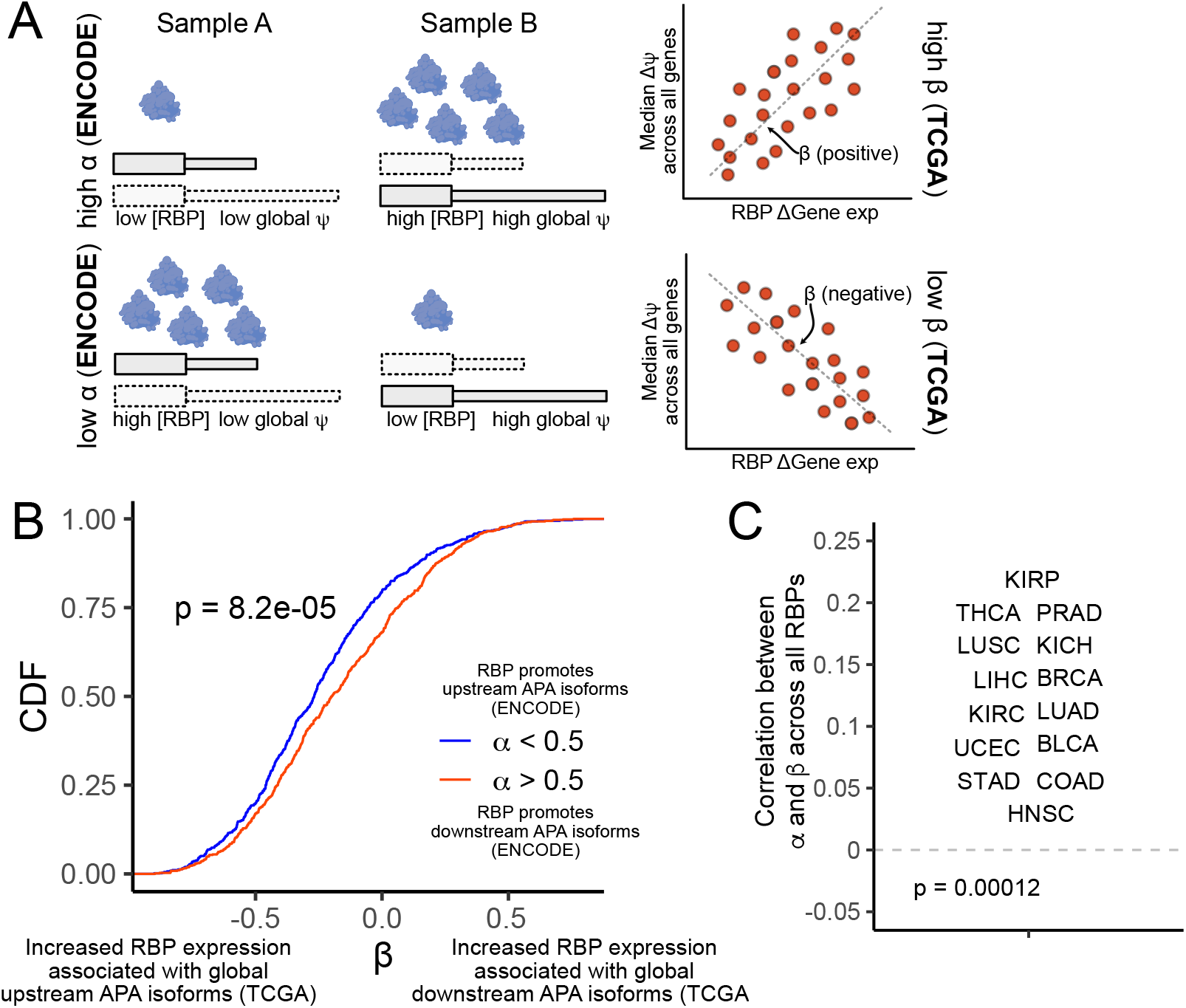
APA is regulated by RBP expression in ENCODE and TCGA data. (A) Diagram depicting connections between changes in RBP expression between condition and widespread, global in changes in *ψ*. Left: In figure 4, RBPs were assigned a value, *α*, based on the effect that their knockdown had on the *Δψ* values for all genes. *α* was defined as the fraction of genes that displayed increased *ψ* values in control knockdown samples compared to RBP knockdown samples. The expression of RBPs with high *α* values was therefore associated with increased *ψ* values transcriptome-wide (top) while expression of RBPs with low *α* values was correlated with decreased *ψ* values transcriptome-wide (bottom). Similar RBP effects were calculated in TCGA data (right) by comparing the change in RBP expression between two matched samples with transcriptome-wide changes in *ψ* values. A value, *β*, was defined as the correlation between changes in RBP expression and the median *Δψ* across all genes with significant *Δψ* values (FDR < 0.05). *α* and *β* are therefore comparable in relating RBP expression and transcriptome wide changes in *ψ* with the former designed for ENCODE RBP knockdown data and the latter designed for TCGA paired sample data. (B) *β* values for RBPs with low *α* values (α < 0.5, blue) and high *α* values (a > 0.5, red). Here, an RBP’s *β* value considers the correlation between its expression and global *ψ* across all TCGA sample pairs. The p value was calculated using a Wilcoxon ranksum test. (C) Correlation between *α* and *β* values across all RBPs for all TCGA sample pairs, separated by cancer type. The p value was calculated using a binomial test for deviation from the expected 0.5 probability that a cancer’s correlation between *α* and *β* would be positive.

If ENCODE-derived effects of RBPs on APA isoform abundance were recapitulated in the TCGA data, we would expect to see a positive correlation between the *α* and *β* values for RBPs. We restricted this comparison to the 94 RBPs that had *α* values significantly different from the expected value of 0.5 (p < 0.01, binomial test). For these RBPs, *α* and *β* values were positively correlated (R = 0.23, p = 0.03). RBPs with *α* values greater than 0.5 had significantly higher *β* values than those with *α* values less than 0.5 (**Figure 7B**). Further, when we correlated *α* and *β* values across all RBPs for all sample pairs within a cancer type, we observed positive correlations in all 12 cancers tested (**Figure 7C**). These results further suggest that dozens of RBPs have the ability to regulate relative APA isoform abundance of many genes in a coordinated, directional manner and that the misregulation of APA seen in many cancers may be due to altered expression of specific RBPs.

## DISCUSSION

Alternative polyadenylation is a key step in control of mRNA function, and its misregulation can have large effects on cellular and even organismal phenotype including major effects on the transcriptome of diseased cells including tumors (Berkovits and Mayr, 2015; Grassi et al., 2018; Mayr and Bartel, 2009; Shi and Manley, 2015; Tian and Manley, 2017; Ulitsky et al., 2012; Zhou et al., 2019). Advances in RNA sequencing and methods of profiling APA from high-throughput data have illuminated the prevalence of APA and its regulation across many cell types and physiological conditions (Ha et al., 2018; Xia et al., 2014). Still, the broad effects of APA on mRNA metabolism, especially beyond changes in mRNA abundance, are not very well understood. Further, the contribution of individual RBPs to the regulation of this process is similarly poorly defined.

To address these challenges, we developed software to accurately quantify alternative polyadenylation and changes in its regulation across conditions from standard RNAseq data. LABRAT builds upon advances in transcriptome quantification using lightweight alignments (Patro et al., 2017) to determine the relative usage of APA sites within genes. This strategy of using fast, accurate, isoform-level quantification has previously been successfully used to study differential isoform regulation (Alamancos et al., 2015; Ha et al., 2018). Here, we have used LABRAT to explore the regulation and consequences of APA in a variety of contexts using thousands of data sets.

The subcellular localization of specific transcripts has been known to be regulated by APA. For example, the dendritic localization of BDNF mRNA depends on the content of the transcript’s 3′UTR as determined by APA (An et al., 2008). More recent transcriptome-wide studies have shown that this phenomenon is widespread, as hundreds of genes display differential enrichments of APA isoforms across cell body and projection compartments (Ciolli Mattioli et al., 2019; Taliaferro et al., 2016; Tushev et al., 2018). Still, there has been confusion as to the relative contributions of tandem UTR- and ALE-mediated APA to this effect, perhaps due to inefficiencies in studying APA with software that uses genomic alignments. LABRAT is the only currently available APA software that explicitly separates and labels these two classes of genes. We took advantage of this to quantify the distribution of tandem UTR and ALE isoforms across subcellular compartments and found that both classes of APA contribute approximately equally to differences in RNA localization. We further found that differential APA isoform localization is most prevalent in young cellular projections that are less than 3 days old, suggesting that this effect may be important for the initiation of projection outgrowth but less significant for the maintenance of established projections.

Although RNA localization is most heavily studied in polarized cell types like neurons, transcripts are asymmetrically distributed in essentially all cells. LABRAT identified hundreds of genes with differential APA isoform enrichment between biochemically defined cytosolic and membrane fractions in nonpolarized D17, HepG2, and K562 cells. These results indicate that APA may play a broad role in subcellular localization to membranes in multiple cell types. The consequences of this localization remain unknown, but given that a large fraction of cellular membrane belongs to the ER, modulation of membrane association may be a way to tune the ER association and therefore translation status of a transcript. Further, given the broad misregulation of APA in many cancers, this may mean that the membrane association of many transcripts changes upon transformation. We further found that genes whose APA isoforms are differentially associated with membranes are less likely to encode ER-targeting signal peptides, suggesting that RNA localization to the ER can occur using mechanisms that are independent of the cotranslational targeting. This phenomenon and its misregulation in specific contexts like cancer needs more study.

The abundance of several CPSF and CstF subunits can have important effects on alternative polyA site choice (Schönemann et al., 2014; Shi et al., 2009; Sun et al., 2018; Takagaki and Manley, 1998). Other RBPs, including CFIm25, have also been shown to strongly directionally regulate APA through activation or repression of specific cleavage events (Masamha et al., 2014; Zhu et al., 2018). Using RBP knockdown followed by high-throughput RNA sequencing experiments performed by the ENCODE consortium (Consortium, ENCODE Project et al., 2012; Davis et al., 2018) we interrogated the regulatory effects of 191 RBPs on APA isoform abundance. In this analysis, the knockdown of dozens of RBPs promoted widespread, coordinated directional shifts in relative APA isoform abundance for hundreds to thousands of genes, suggesting that the repertoire of RBPs that can differentially regulate APA isoforms is quite large. It is important to note, though, that not all of these RBPs may be directly regulating APA. For example, many may be differentially regulating stability of 3′ UTR isoforms.

The CPA apparatus processes nascent Pol II transcripts at the ends of genes in the context of complexes with Pol II. According to the “window of opportunity” model (Bentley, 2014), the decision between alternative polyA sites can be influenced by the delay between synthesis of upstream and downstream sites which is determined by the speed of transcription. Consistent with this model, we found using LABRAT that slow transcription caused by a mutation in the Pol II large subunit causes a significant shift in favor of upstream polyA sites and that this effect is true for both the ALE and tandem 3’ UTR classes of APA. Moreover, as predicted by the “window of opportunity” model the mRNAs with the greatest upstream shift in APA correspond to those with the greatest distance between alternative tandem 3’UTR sites (**Figure 3D**). In summary, these results show that Pol II speed can significantly modulate alternative polyA site choice. They further suggest the possibility that regulation of transcription elongation could contribute to changes in APA under normal and pathological conditions.

Connections between APA and cancer have been well established (Masamha and Wagner, 2018; Masamha et al., 2014; Xia et al., 2014). Generally, conclusions regarding this relationship have been focused on the idea of increased proximal APA in cancerous samples (Masamha et al., 2014; Mayr and Bartel, 2009; Xia et al., 2014) with the idea that proximal APA of oncogenic transcripts particular removes repressive regulatory elements in the distal UTR that might otherwise keep the expression of these genes low. However, our results using LABRAT to assess APA changes in 671 paired tumor and normal samples indicate that broad, directional shifts in APA are specific to the type of cancer being studied. Some cancers, including lung cancers and head-neck squamous cell carcinoma, display the canonical increased use of proximal APA sites, while others, including kidney renal clear cell carcinoma and thyroid cancers, show strong shifts in the opposite direction toward distal APA sites. Further, increased proximal and distal APA is associated with poor patient prognosis in head-neck squamous cell carcinoma (HNSC) and kidney renal clear cell carcinoma (KIRC), respectively. Critically, this indicates that increased proximal APA is not a general signature of cancer, but rather that the direction of APA misregulation is cancer-specific.

Relationships between APA and gene expression have also been well documented (Mayr and Bartel, 2009; Sandberg et al., 2008). Early studies of this connection indicated that distal APA was generally associated with a decrease in gene expression. Later studies, though, indicated that this relationship was less clear (Spies et al., 2013). To investigate how APA affects gene expression, we compared changes in *ψ* values and changes in gene expression for all genes in over 1000 pairs of RNAseq samples. We found that within a sample, correlations between gene expression and APA were weak, but were consistently in the canonical, expected direction where distal APA leads to lower expression. Reorienting the analysis to interrogate the relationship within single genes but across samples again revealed that the average gene has only a very weak connection between APA and gene expression. Still, some genes had remarkable correlations (R ~ 0.7-0.8) between these two measurements, indicating that changes in their expression across diverse samples are controlled in large part by modulation of APA site choice.

Across over a thousand pairs of samples, we observed strong correlations between APA changes in genes with tandem UTRs and those with ALEs. If a particular condition promoted increased distal APA in tandem UTR genes, it overwhelmingly also promoted increased distal APA in ALE genes and vice versa. This strongly indicates that the two may be regulated by similar mechanisms. Tandem UTRs are regulated solely at the level of cleavage/ polyadenylation. The simplest interpretation of our results is therefore that the contribution of regulated splicing to ALE control is minor compared to that of regulated cleavage/polyadenylation, perhaps because splicing kinetics are slower. For ALEs, proximal cleavage events obviate potential regulation of the ALE by splicing since the distal ALE is removed from the transcript. If recognition of the proximal APA site by the cleavage and polyadenylation machinery is inhibited, this may provide time for splicing to distal ALEs to occur, and this decision could be affected by the speed of transcription. In this model, splicing acts on ALEs only if given the chance to do so through inhibition of kinetically favored cleavage events.

Overall, the results presented here shed light on the molecular consequences of APA and make predictions about the proteins and mechanisms involved in its regulation. Further experimental studies are needed to fully understand these processes. We envision LABRAT as an important tool in deriving meaningful insights from those experiments.

## ACKNOWLEDGEMENTS

We thank Neel Mukherjee and members of the Taliaferro Lab for helpful discussions regarding analyses. We also thank Rob Patro for helpful discussions regarding using Salmon-based transcript quantification for APA investigation.

The Genotype-Tissue Expression (GTEx) Project was supported by the Common Fund of the Office of the Director of the National Institutes of Health, and by NCI, NHGRI, NHLBI, NIDA, NIMH, and NINDS.

This work was funded by the National Institutes of Health (R35-GM133885 to J.M.T. and R35-GM118051 to D.B.) and the RNA Bioscience Initiative at the University of Colorado Anschutz Medical Campus (J.M.T.). It was further supported by a Predoctoral Training Grant in Molecular Biology (NIH-T32-GM008730) (R.G.).

## METHODS

### General LABRAT usage

LABRAT is freely available for download here: https://github.com/TaliaferroLab/LABRAT/. LABRAT searches for specific tags in the annotation associated with transcripts with ill-defined 3’ ends. These tags are present in Gencode (www.gencodegenes.org) gff annotations but may not be present in annotations from other sources. For this reason, we strongly suggest using Gencode annotations for use with LABRAT. For analysis of Drosophila data, we modified LABRAT to perform similar filtering on Ensembl annotations for the dm6 Drosophila genome build. This version of LABRAT is also available at the above GitHub address.

Genes that did not pass an expression filter (TPM ≥ 5) were removed from further analysis. This gene expression was defined as the sum of the expression values for all valid, filter-passing transcripts for the gene. LABRAT reports these genes as having a *ψ* value of NA.

Identification of genes with significantly different *ψ* values across conditions was done using a linear mixed effects model with the Python package statsmodels (Seabold and Perktold, 2010). For simple comparisons involving two conditions, a simple model relating conditions and *ψ* values was used (*ψ* values ~ condition). For analysis of the CeFra and ENCODE data, slightly more complex models were used. In the CeFra data, the method of library preparation, polyA-enrichment or ribosomal RNA depletion, was added as a covariate (*ψ* values ~ condition + libprep). In the ENCODE data, the cell line, K562 or HepG2, was added as a covariate (*ψ* values ~ condition + cell line). These models were then compared to null models where the effect of the condition was removed. For simple comparisons, the null models were specified as (*ψ* values ~ 1). For the CeFra and ENCODE comparisons, these were specified as (*ψ* values ~ libprep) and (*ψ* values ~ cell line), respectively. A likelihood ratio test was then used to evaluate the relative fit between the experimental and null models. P values were derived from the likelihood ratio test and then corrected for multiple hypothesis testing using a Benjamini-Hochberg correction (Benjamini and Hochberg, 1995). *Δψ* values are defined as differences in mean *ψ* across conditions.

To define tandem UTR and ALE structures, LABRAT observes the isoform structures at the 3’ end of a gene. If all APA sites are contained within the same exon, then the structure in tandem UTR. If all APA sites are contained within different exons, then the structure is ALE. If a gene has only two APA sites, then its structure must be either tandem UTR or ALE. If a gene has more than two APA sites, it is possible for the gene to fit into neither classification. For example, in a gene with three APA sites, it is possible to have two of them contained within one exon and the third by itself in another exon. In these cases, LABRAT assigns the gene to have a “mixed” structure.

### Comparison of APA in mouse brain and liver tissues

RNAseq data for mouse brain and liver tissues was downloaded from (https://www.ncbi.nlm.nih.gov/bioproject/?term=PRJNA375882) (Li et al., 2017). Each tissue sample contained 8 replicates. Genes with significantly different *ψ* values were identified as those with an FDR of less than 0.05.

### Analysis of APA in GTEx RNAseq data

RNAseq data from the Genotype-Tissue Expression (GTEx) project (BioProject PRJNA75899) were downloaded from the NCBI Sequence Read Archive (SRA) via dbGaP-authenticated access and quantified using salmon (Patro et al., 2017) as described elsewhere in this manuscript. *ψ* values were calculated for each gene in each sample using LABRAT. LABRAT employs an expression level cutoff, returning a *ψ* value of NA if the sum of expression of all isoforms for a gene is not at least 5 TPM. There were many genes in this analysis of tissue-specific RNAseq that therefore had *ψ* values of NA in at least one sample. To facilitate PCA analysis, these missing *ψ* values were imputed using the R package missMDA (Josse and Husson, 2016).

The data used for the analyses described in this manuscript were obtained from dbGaP accession number phs000424.vN.pN between 07/16/2020 and 08/31/2020.

### APA analysis of simulated RNAseq data

To compare the performance of LABRAT to QAPA (Ha et al., 2018), DaPars (Xia et al., 2014) and Roar (Grassi et al., 2016), we generated a synthetic RNAseq dataset. In this dataset, 5000 genes with only two alternative polyadenylation sites were analyzed. 1250 were randomly assigned to have positive *Δψ* values, 1250 were assigned to have negative *Δψ* values, and 2500 were assigned to have no significant change in *ψ* between conditions. Each gene was then randomly assigned a TPM expression value using a Dirichlet distribution with numpy.random.dirichlet.

The simulation was performed by comparing three replicates each from two conditions. For the positive *Δψ* genes, the minimum *ψ* from condition B was required to be at least 0.1 greater than the maximum *ψ* from condition A. Conversely, for the negative *Δψ* genes, the maximum *ψ* from condition B was required to be at least 0.1 less than the minimum *ψ* from condition A. For control genes, the difference between any two *ψ* values both within and across conditions was required to be less than 0.25.

Given a gene’s overall expression and its *ψ* value, TPM values were then relatively split between polyadenylation sites such that the desired *ψ* value was achieved. TPM values for individual transcripts within polyadenylation sites were then assigned. If a polyadenylation site was only supported by a single transcript, that transcript was given the site’s entire TPM value. If a polyadenylation site was supported by multiple transcripts, the site’s TPM allotment was randomly distributed among the transcripts.

Given a transcript’s assigned TPM value and its length, the desired number of counts for each transcript was then computed by multiplying the TPM value by the length of the transcript. The sequence of each transcript and the desired number of counts were then given to the R package polyester (Frazee et al., 2015) to create synthetic, 100 nucleotide, paired-end RNAseq reads.

In analyzing the reads with each package, gene assignments (positive *Δψ*, negative *Δψ*, or control) made by the software were compared to the assignments made during preparation of the synthetic dataset. For analysis of these reads with LABRAT, genes with FDR values of less than 0.05 were called as affected genes (either positive or negative *Δψ* depending on the reported *Δψ* value) while those with values of 0.05 or greater were called as control genes. For analysis with QAPA, genes with differences in PPAU values of at least 10 were called as affected genes while those with differences in PPAU values of less than 10 were called as control genes. For analysis with DaPars, genes with adjusted p values of less than 0.05 were called as affected genes while those with adjusted p values of 0.05 or greater were called as control genes. For analysis with Roar, genes with p values less than 0.05 and roar values greater than 1.1 were called as positive *Δψ* genes, genes with p values less than 0.05 and roar values less than 0.9 were called as negative *Δψ* genes, while genes with p values of 0.05 or greater were called as control genes.

### Analysis of differential APA isoform enrichment across subcellular compartments

*ψ* values for each subcellular compartment were quantified using LABRAT, and genes with significant changes in *ψ* values across compartments were identified using an FDR cutoff of 0.05. The fraction of these significant genes with greater *ψ* values in the projections than cell bodies was calculated. Binomial p values were calculated for deviations from the expected fraction of 50%. Times of projection growth were manually curated from the methods description of each study.

### Analysis of differential APA isoform enrichment across biochemically defined subcellular fractions

*ψ* values for each subcellular fraction were quantified using LABRAT, and genes with significant changes in *ψ* values across compartments were identified using an FDR cutoff of 0.05. FDRs were calculated using a linear model that incorporated the method of library preparation (polyA-enrichment or ribosomal RNA depletion) as a covariate.

### Quantification of ER signal sequence abundance

For each gene, the translation of its longest CDS sequence was given to the signal sequence prediction program SignalP (Almagro Armenteros et al., 2019). For a set of genes, the fraction of genes within the set that contained at least one SignalP-defined ER signal sequence was calculated. For comparing these fractions across sets of genes, a distribution of fractions was created by bootstrapping where 40% of the genes were sampled 100 times.

### Analysis of APA changes induced by changes in RNA Polymerase II speed

RNAseq data from HEK293 cells expressing slow (R749H) and wildtype RNA polymerase II (Fong et al., 2014) were downloaded from the Gene Expression Omnibus (GSE63375). Using an FDR cutoff of 0.1, genes with significantly different *ψ* values between wildtype and R749H samples were identified using LABRAT.

### Analysis of ENCODE RBP RNAi knockdown RNAseq samples

In this dataset, each RBP was associated with two RBP RNAi samples and two control RNAi samples. We limited analyses to RBPs that had knockdown samples in both K562 and HepG2 cell lines. *ψ* values were calculated comparing RBP knockdown and control knockdown samples, and genes with significant *ψ* differences between RBP RNAi and control RNAi samples were identified using an FDR cutoff of 0.05. FDRs were calculated using a linear model that incorporated the cell line (HepG2 or K562) as a covariate.

For each RBP, the fraction of these significant genes with greater *ψ* values in the control RNAi than RBP RNAi was calculated. These fractions were defined as a value, *α*, where *α* ranged from 0 to 1. *α* values greater than 0.5 were therefore associated with larger *ψ* values (and therefore more distal APA) in the control RNAi sample. Conversely, *α* values less than 0.5 were therefore associated with smaller *ψ* values (and therefore more proximal APA) in the control RNAi sample. Each RBP was therefore assigned one *α* value from the ENCODE data. Binomial p values were calculated for deviations from the expected fraction of 50%.

### Comparison of ENCODE RBP RNAi knockdowns and eCLIP RBP binding data

The eCLIP narrowpeak bed files for isogenic replicates aligned to GRCh38 for each RBP measured in both HepG2 (103 RBPs) and K562 (120 RBPs) were downloaded from www.encodeproject.org. Analyses were restricted for within each line and not combined. For each individual RBP data set, overlapping peaks were merged using bedtools v2.29.2 (Quinlan and Hall, 2010). These peaks were then intersected with the longest 3’UTR of genes whose polyA sites were both affected and unaffected by RBP knockdown (as measured by LABRAT described above). RBP occupancy was scored for each 3’UTR as either present or not. The statistical significance of a given RBPs occupancy within the subset of genes whose polyA site choice was affected by knockdown of any RBP was determined using a binomial test.

The number of RBPs that were ‘self significant’, i.e. the occupancy of a specific RBP was significant for the genes whose polyA site choice was affected by knockdown of that same RBP, was determined for both HepG2 and K562. To determine if that number was greater than what was expected by chance, relationships between RBPs and the genes they bind were shuffled, and the analysis was repeated to identify the number of ‘self significant’ RBPs. This process was repeated 1000 times to generate a null distribution of the number of ‘self significant’ RBPs. The number of actual ‘self significant’ RBPs was then compared to the null distribution and an empirical p value was calculated.

### Analysis of APA in TCGA matched tumor/normal tissue samples

In this dataset, each patient is associated with a pair of samples, one from a tumor and another from matched normal tissue. *ψ* values were calculated for each sample, and genes with significant *ψ* differences between all tumor samples and all normal samples within a cancer type were identified using an FDR cutoff of 0.05.

Using the TCGA data, the effect of an RBP’s expression on *ψ* was inferred by correlating changes in the RBP’s expression across samples with changes in *ψ* values of genes that passed the FDR cutoff of 0.05. For each tumor/normal pair, the change in RBP expression was calculated by comparing TPM expression values, and changes in *ψ* were calculated by finding the median *Δψ* value across genes with significant changes in *ψ*. The spearman correlation coefficient of this comparison across all tumor/normal pairs was defined as *β*. Each RBP was therefore assigned one *β* value from the TCGA data.

### Analysis of survival data in TCGA samples

Using the tumor and matched normal tissue samples from the TCGA dataset, genes with significant *ψ* differences (FDR < 0.05) were identified for each tumor type as misregulated genes. The median *ψ* of misregulated genes was then calculated for each patient in samples without matched normal tissue controls. Patients were then ranked by their median *ψ* of misregulated genes and separated into quartiles. Only patients within the most extreme quartiles were plotted for each tumor type.

Clinical data for each patient was obtained from cbioportal (Gao et al., 2013). Survival analysis and plotting was performed with R packages survival (version=3.1-8) (Therneau and Grambsch, 2000) and survminer (version=0.4.8) (Alboukadel Kassambara, 2020). Log-rank tests for significance were calculated to compare extreme quartiles for each tumor type and were considered significant if less than 0.05.

### Analysis of relationship between APA and RNA expression

For every pair of samples (Control and RBP RNAi in ENCODE and tumor/normal samples in TCGA), the change in RNA expression and *ψ* value for every gene was calculated. Gene expression filters (TPM ≥ 5) were applied, but FDR cutoffs for *Δψ* were not. These two values were then compared to each other, and the resulting Spearman correlation coefficient was defined as rho (*ρ*). If distal APA (i.e. increases in *ψ*) was associated with decreases in RNA expression, the resulting *ρ* value would be negative.

*ρ* was calculated in two different ways. In the first way, changes in expression and *ψ* for all genes within a sample were correlated. In this comparison, each sample pair ends up with a single *ρ* value. In the second way, changes in expression and *ψ* for a single gene across all sample pairs were correlated. In this comparison, each gene ends up with a single *ρ* value in each sample set (ENCODE and TCGA).

The second *ρ* calculations were used to categorize genes as being either positively or negatively correlated. To achieve similar numbers of genes in each category, a positive *ρ* in either sample set was considered as positively correlated while a *ρ* less than −0.15 in either sample set was considered negatively correlated. Genes behaving inconsistently between sample sets were removed from these categories and placed in the control gene category(25% of positively correlated and 14% or negatively correlated). For simplicity, genes with only two APA isoforms were considered during this categorization resulting in 316 positively correlated genes, 313 negatively correlated genes and 1466 control genes used in UTR sequence analysis.

### Quantifying effects on RNA expression due to UTR content with qRT-PCR

Proximal and distal UTR regions were cloned onto the coding sequence of Firefly luciferase. In this plasmid, Firefly luciferase is driven by a bidirectional tet-On promoter. This promoter also drives Renilla luciferase, which served as a control in these experiments. The resulting plasmids were transfected into HeLa cells using Lipofectamine 2000 (Life Technologies). These cells were engineered to contain a single loxP-flanked cassette within their genome (Khandelia et al., 2011). The plasmid was site-specifically integrated into the genome of the HeLa cells by cotransfecting it with a plasmid expressing Cre recombinase. Recombinants were then selected using 1 μg / mL puromycin for 2 weeks.

The expression of Firefly and Renilla luciferase transcripts was induced by incubating cells with 1 μg / mL doxycycline for 48 hours. Total RNA was then isolated using a Quick RNA Isolation Mini Kit (Zymo Research). 1 μg of total RNA was reverse transcribed using iScript Reverse Transcriptase Supermix (BioRad). The relative levels of Firefly and Renilla luciferase transcripts in the sample were then quantified using Taqman qPCR. For each gene, the ratio of Firefly to Renilla luciferase in the case where the proximal UTR was fused to Firefly luciferase was set to 1.

### Identifying features enriched in UTRs associated with gene expression changes

For each gene considered in this analysis (positively correlated, negatively correlated and control genes), proximal and distal UTR sequences were extracted in such a way that they contained unique sequences only. This means that the distal UTRs of genes with tandem UTR models lacked the beginning of their sequence which is unique to the proximal UTR as illustrated in Figure 6H.

UTR sequence features of either positively or negatively correlated genes were always compared to the control gene set. Enrichment analyses were performed using a custom R package (FeatureReachR) publicly available here: https://github.com/TaliaferroLab/FeatureReachR. This R package utilizes wilcoxon ranksum tests to compare length and GC contents of the three gene sets. Motif and five-mer enrichment significance is calculated with a Fisher’s exact test and corrected using the Benjamini & Hochberg method (Benjamini and Hochberg, 1995). RBP binding motifs are represented as a sequence match > 80% with position weight matrices sourced from the CISBP-RNA database (http://cisbp-rna.ccbr.utoronto.ca/) (Ray et al., 2013) or RNA bind-N-seq results (Dominguez et al., 2018). AREScore (Spasic et al., 2012) was utilized to determine the presence of AU rich elements within the UTRs and compared again using wilcoxon rank-sum tests (http://arescore.dkfz.de/arescore.pl).

## SUPPLEMENTARY FIGURES

**Figure S1.**
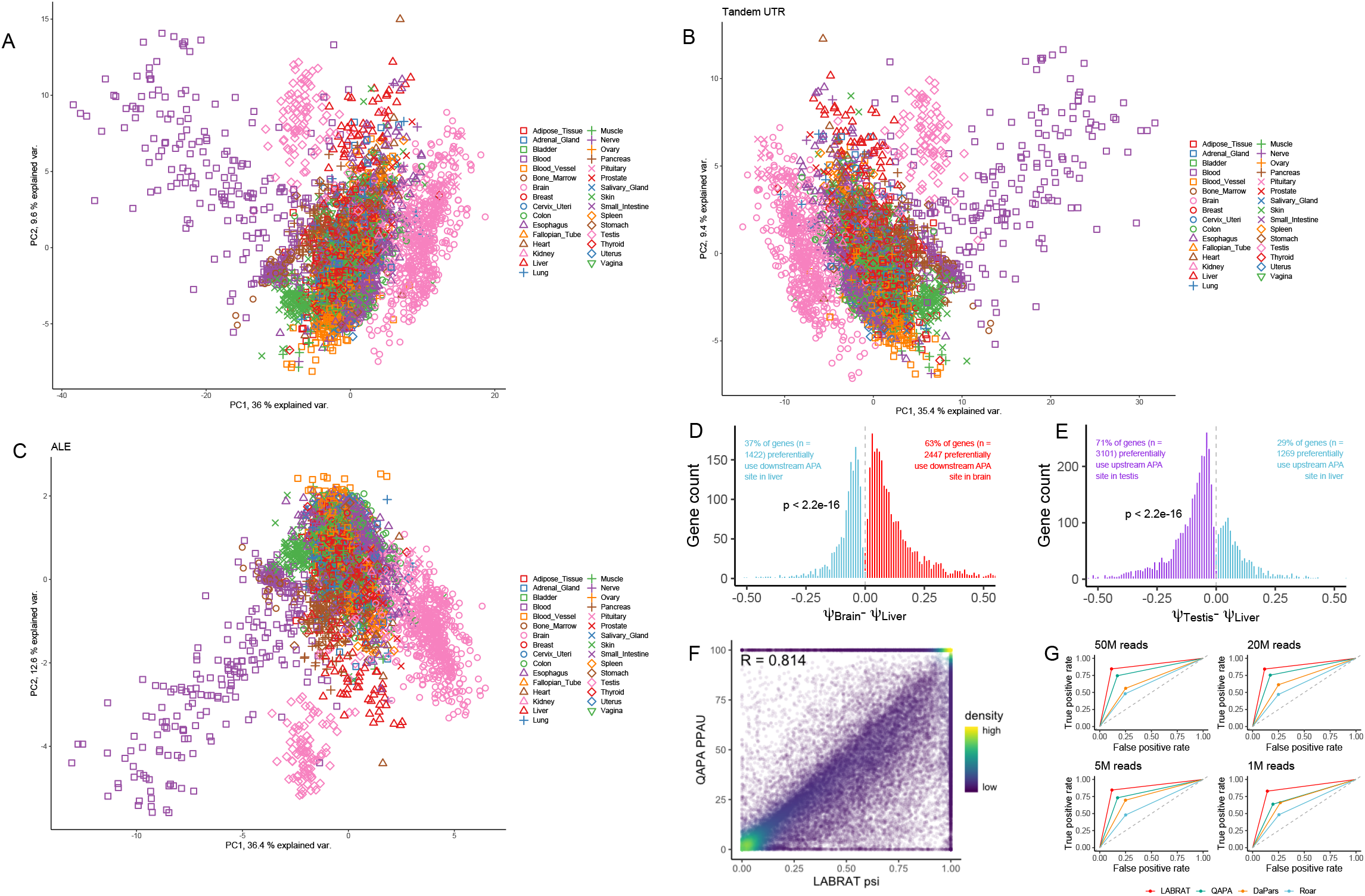
(A) PCA analysis of *ψ* values calculated from human tissues. Data was produced as part of the GTEx project. (B) As in A, but only using genes that have a tandem UTR APA structure. (C) As in A, but using only genes that have an ALE APA structure. (D) Comparison of *ψ* values from human brain and liver samples. Delta *ψ* values for genes with FDR values less than 0.01 are plotted. (E) Comparison of *ψ* values from human testis and liver samples. Delta *ψ* values for genes with FDR values less than 0.01 are plotted. (F) Comparison of APA quantifications produced by LABRAT (ψ) and QAPA (PPAU). (G) Benchmarking of APA software performance at a range of sequence read depths.

**Figure S2.**
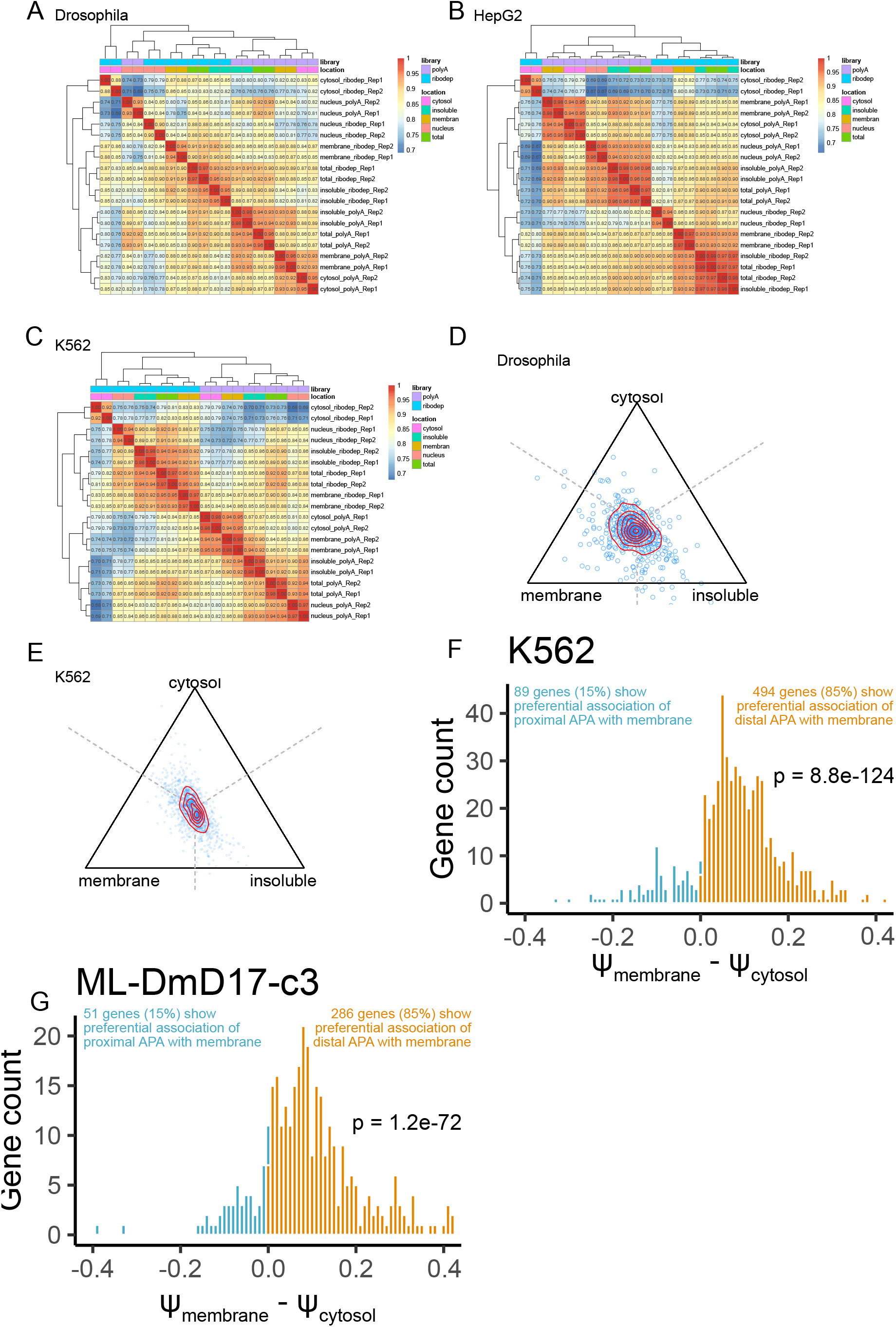
Hierarchical clustering of *ψ* values from biochemically fractionated *Drosophila* DM-D17-C3 cells (A), HepG2 cells (B), and K562 cells (C). (D-E) Simplex plots relating relative *ψ* values for genes between the cytosolic, membrane-associated, and insoluble fractions of DM-D17-C3 cells (D) and K562 cells (E). A dot that is equidistant from all three vertices had equal *ψ* values in each fraction while a dot that is closer to one vertex had a higher *ψ* value in that fraction relative to the other two fractions. (F-G) Comparison of *ψ* values in K562 (F) and DM-D17-C3 (G) cytosolic and membrane fractions for genes whose *ψ* value was significantly different between these compartments (FDR < 0.01).

**Figure S3.**
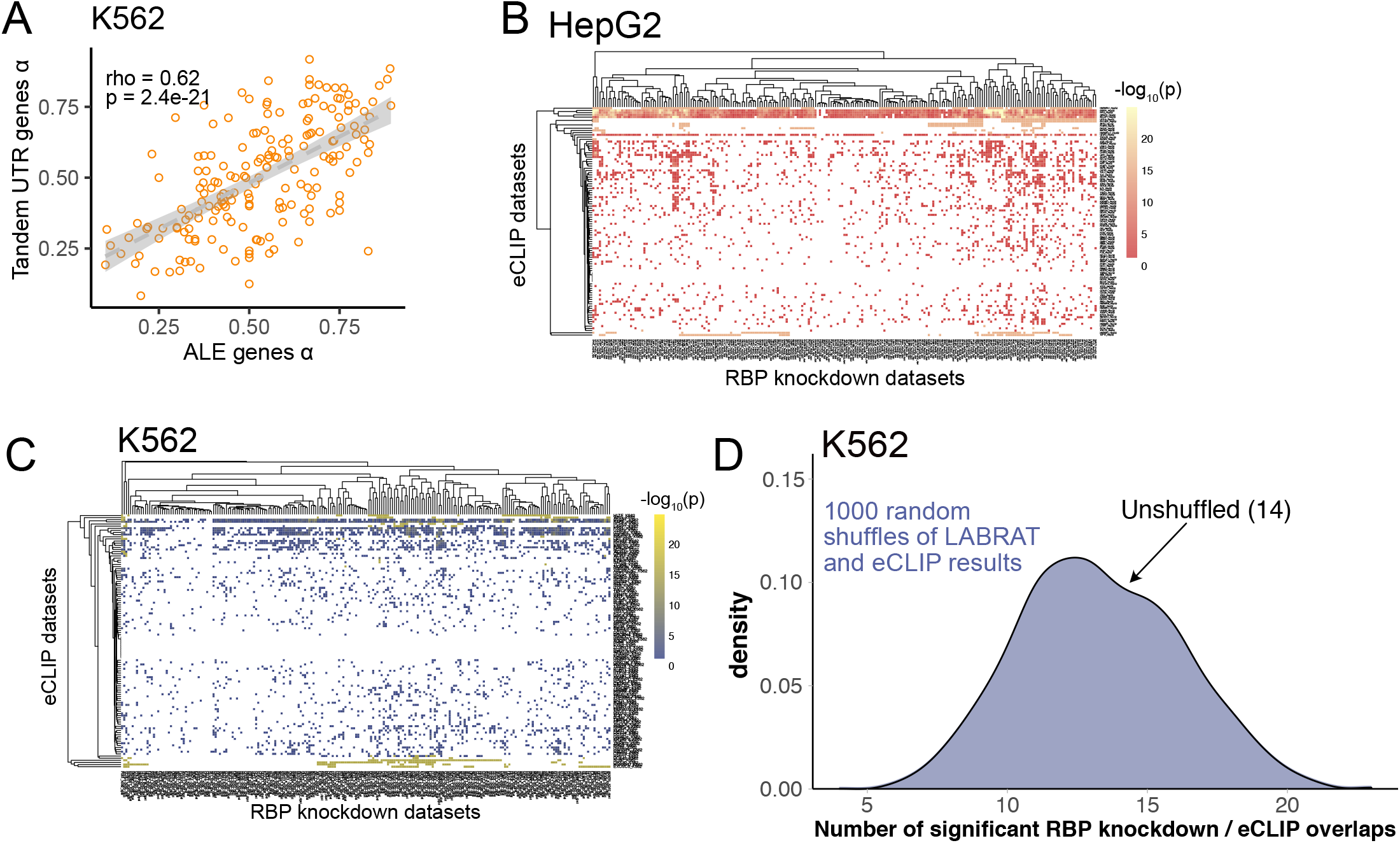
(A) *α* values for each RBP knockdown in K562 cells were calculated using tandem UTR and ALE genes independently. These were then plotted and correlated. Each dot in this plot represents one RBP knockdown experiment. (B) Binomial p values for overlaps between genes whose APA was sensitive to RBP knockdown and genes whose 3’ UTRs were bound by an RBP in eCLIP experiments. Data taken from ENCODE HepG2 experiments. (C) As in B, but using data from ENCODE K562 experiments. (D) As in Figure 4E. Among 102 RBPs expressed in K562 cells, overlaps between the genes whose APA was sensitive to RBP knockdown and the genes whose 3’ UTRs were bound by the RBP in eCLIP experiments were calculated. The significance of this overlap was calculated using a binomial test. 14 RBPs bound the 3’ UTRs of their APA targets more often than expected (binomial p < 0.05). To assess whether this was more than the expected number of significant RBPs, relationships between RBPs and their lists of APA and eCLIP targets were shuffled 1000 times, and the analysis was repeated after each shuffle to create a null distribution (blue).

**Figure S4.**
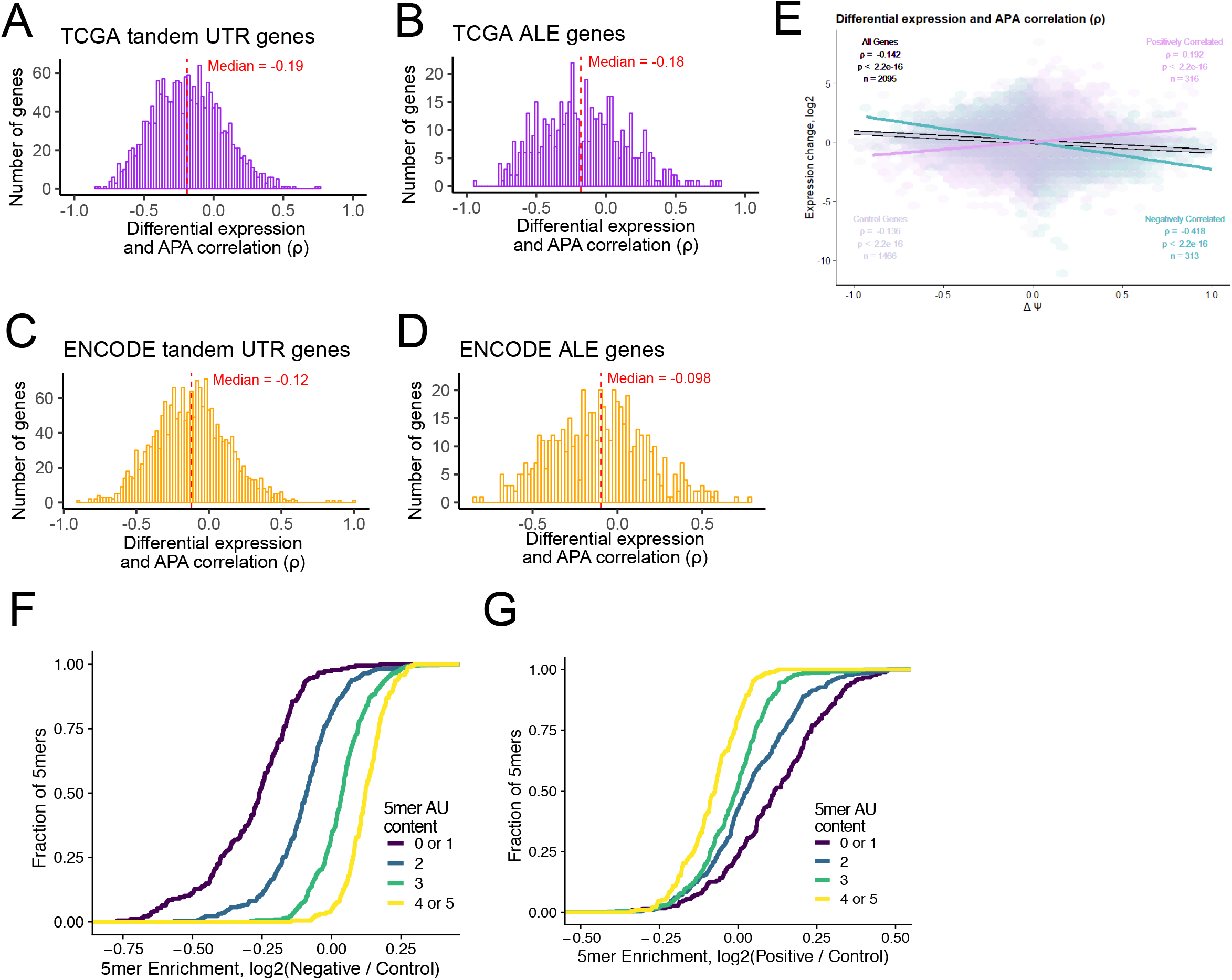
(A-B) Histogram of gene-wise correlations between changes in ≠ and changes in gene expression (o) derived from TCGA tumor and matched normal samples for tandem UTR (A) genes and ALE (B) genes. (C-D) Histogram of gene-wise correlations between changes in ≠ and changes in gene expression (*ρ*) derived from ENCODE RBP knockdown and control samples for tandem UTR (C) genes and ALE (D) genes. (E) Binned scatter plot comparing changes in ≠ and changes in gene expression for genes with negative *ρ* values (blue), positive *ρ* values (purple) and control genes (gray). (F) Enrichment of 5mers in the distal UTRs of negatively correlated genes compared to the distal UTRs of control genes. (G) Enrichment of 5mers in the distal UTRs of positively correlated genes compared to the distal UTRs of control genes.

